# snoFlake: A network model for snoRNA–RBP complexes reveals SNORD22 as a U5 snRNP-associated splicing regulator

**DOI:** 10.64898/2026.04.02.716167

**Authors:** Kristina Sungeun Song, Marianne Cyr, Laurence Faucher-Giguère, Bernice Yeo, Vivian Kaiyan Seow, Gabrielle Deschamps-Francoeur, Sherif Abou Elela, Michelle S. Scott

**Affiliations:** Département de biochimie et de génomique fonctionnelle, Faculté de médecine et des sciences de la santé, Université de Sherbrooke, Sherbrooke, Québec, J1E 4K8, Canada; Département de microbiologie et d’infectiologie, Faculté de médecine et des sciences de la santé, Université de Sherbrooke, Sherbrooke, Québec, J1E 4K8, Canada

**Keywords:** snoRNA–RBP interactions, interaction network, alternative splicing, U5 snRNP, post-transcriptional regulation

## Abstract

Small nucleolar RNAs (snoRNAs) are canonically viewed as stable components of ribonucleoprotein complexes dedicated to RNA modification. Here, we developed snoFlake, a snoRNA-centric interaction network integrating physical and functional associations between box C/D snoRNAs and RNA-binding proteins (RBPs), challenging this narrow view. Using snoFlake, we systematically identified snoRNAs predicted to form noncanonical complexes with diverse RBPs, extending their roles into post-transcriptional regulation. We found 23 high-confidence network motifs enriched for RNA-processing functions, including a top-ranked module linking SNORD22 to U5 snRNP components PRPF8 and EFTUD2. SNORD22 co-binds with these spliceosomal RBPs at splice sites showing reduced U5 snRNP occupancy, suggesting a role in reinforcing spliceosomal engagement at suboptimal exons. Consistently, SNORD22 depletion promotes exclusion of weak cassette exons, altering transcript isoform composition and predicted coding output. Beyond SNORD22, snoFlake reveals snoRNAs with similar network profiles, providing a resource for uncovering previously uncharacterized snoRNA–RBP complexes and expanding the functional snoRNome.

## Introduction

Small nucleolar RNAs (snoRNAs) are a conserved and abundant class of noncoding RNAs that primarily function as guides for RNA modification. They associate with specific sets of proteins to form small nucleolar ribonucleoprotein complexes (snoRNPs), which mediate site-specific modifications on target RNAs^1^. These modifications play a central role in the maturation of ribosomal RNAs (rRNAs) and small nuclear RNAs (snRNAs), thereby facilitating the assembly and function of the ribosome and spliceosome, respectively^2–4^.

SnoRNAs are broadly classified into two major families based on conserved sequence motifs and associated catalytic activities. Box C/D snoRNAs guide 2′-O-methylation by recruiting the methyltransferase fibrillarin (FBL), whereas box H/ACA snoRNAs direct pseudouridylation through association with the pseudouridine synthase dyskerin (DKC1)^5,6^. In addition to these catalytic subunits, each class forms a stable snoRNP with a conserved set of core proteins that ensure proper folding, localization, and protection of the snoRNA. Box C/D snoRNP components include NOP56, NOP58, and the kink-turn-binding protein SNU13, while H/ACA snoRNPs associate with GAR1, NHP2, and NOP10^7^. Target recognition is typically mediated through short antisense elements (ASEs) within the snoRNA that base-pair with complementary sequences in substrate RNAs, enabling modification with high specificity^5^.

Despite this well-defined canonical framework, a substantial fraction of human snoRNAs lack known modification sites on rRNAs or snRNAs^8,9^. These so-called orphan snoRNAs have challenged the traditional view of snoRNA function and prompted investigations into alternative regulatory roles. Indeed, recent studies have expanded the repertoire of snoRNA-guided modifications to include noncanonical targets such as transfer RNAs (tRNAs) and messenger RNAs (mRNAs), implicating snoRNAs in the regulation of tRNA biogenesis and stability^10^ as well as mRNA translation through site-specific modification^11^. These findings suggest that snoRNAs participate in broader layers of post-transcriptional regulation beyond ribosome and spliceosome biogenesis.

Accumulating evidence further indicates that many noncanonical snoRNA functions are mediated through interactions with RNA-binding proteins (RBPs) beyond the core snoRNP components. Certain snoRNAs act as molecular scaffolds or decoys that influence RNP complex assembly. For example, the H/ACA snoRNA SNORA73 facilitates the formation of a complex between 7SL RNA and mRNAs encoding proteins of the secretory pathway^12^. Other snoRNAs directly regulate protein activity: SNORD50A and SNORD50B interact with Ras family GTPases to inhibit Ras farnesylation and membrane localization^13^, while SNORD50A has also been shown to bind the cleavage and polyadenylation factor FIP1 to influence 3′-end processing^14^. Additional roles in RNA metabolism have been described, including the recruitment of the TRAMP–exosome complex by SNORD63-derived fragments to promote selective nuclear RNA decay^15^. Collectively, these examples point to a recurring theme in which snoRNAs engage diverse RBPs to exert regulatory functions distinct from their canonical modification activities.

One particularly compelling yet poorly understood area concerns the potential involvement of snoRNAs in pre-mRNA splicing. Several box C/D snoRNAs have been proposed to modulate alternative splicing by base-pairing with regulatory elements in pre-mRNAs. SNORD27, for instance, binds the *E2F7* transcript and modulates exon inclusion independent of its canonical methylation function^16^. Similarly, SNORD115 has long been associated with splicing regulation of the serotonin receptor 2C mRNA, with more recent evidence suggesting broader, sequence-specific effects across the transcriptome^17,18^. Additional studies have reported interactions between snoRNAs and splicing factors, including PTBP1 and SMNDC1^19^. However, the scope, organization, and mechanistic basis of snoRNA–spliceosome interactions remain unresolved. In particular, it is unclear whether snoRNAs can act *in trans* to guide spliceosomal activity through sequence-specific RNA interactions, analogous to their canonical role in RNA modification.

A major limitation in addressing these questions has been the lack of a systematic framework to map snoRNA interactions with RBPs and RNA targets at scale. For decades, the snoRNA interactome has been largely defined by the core proteins required for snoRNP assembly and catalytic function^7^. While isolated studies have revealed individual snoRNA–RBP interactions with regulatory potential, how these interactions are organized, whether they form coherent functional modules, and how they contribute to post-transcriptional regulation at the systems level remain unexplored^1,8^.

To address this gap, we developed snoFlake (<u>sno</u>RNA <u>F</u>unctional Inter<u>a</u>ction Networ<u>k</u> Mod<u>e</u>l), an integrative interaction network that maps physical and functional connections between human box C/D snoRNAs and RBPs. By integrating high-throughput and computationally predicted interaction datasets, snoFlake uncovers thousands of high-confidence snoRNA–RBP associations, many of which involve canonical guide snoRNAs and splicing-related RBPs. We introduce “double-edge” interactions, defined as snoRNA–RBP pairs supported by both physical binding and co-binding on shared protein-coding RNA targets. Using these interactions as building blocks for network motif discovery, we identified 23 snoRNA-centered motifs significantly enriched beyond random expectation. These motifs were frequently composed of RBPs with shared functional annotations, suggesting cooperative snoRNA–RBP assemblies with specialized regulatory capacities.

One such example is SNORD22, a highly expressed orphan box C/D snoRNA^20^ that forms a tripartite complex with U5 snRNP components PRPF8 and EFTUD2. Together, they co-bind a subset of weakly spliced cassette exons at splice-site-proximal regions where basal PRPF8 and EFTUD2 occupancy is comparatively reduced. SNORD22 promotes inclusion of these exons, supporting a compensatory, *trans*-acting role in facilitating U5 snRNP recruitment within suboptimal splicing contexts. Beyond SNORD22, snoFlake uncovers additional box C/D snoRNAs with similar interaction architectures, highlighting its utility as a systematic framework for mapping noncanonical snoRNA–RBP complexes and their roles in post-transcriptional gene regulation.

## Results

### snoFlake: A functional interactome of human box C/D snoRNAs and RNA-binding proteins

To systematically investigate the landscape of human box C/D snoRNA-protein interactions, we developed snoFlake, a heterogeneous network that captures both physical and functional relationships between snoRNAs and RBPs (**Figures 1A and S1**). In this network, snoRNAs and RBPs are represented as nodes, with edges denoting physical binding or functional co-association on shared protein-coding RNA targets. We built snoFlake using a reproducible Snakemake-based^21^ pipeline with curated inputs: annotated human box C/D snoRNAs from snoDB (v2.0)^20^ and RBPs profiled by eCLIP in the ENCODE project^22^. To ensure biological relevance and detectable expression, both snoRNAs and RBPs were filtered by transcript abundance, defined as ≥ 1 transcript per million (TPM) in at least one TGIRT-seq dataset (see **STAR Methods**), a method that provides accurate quantification of both structured and unstructured RNAs^23,24^. Additional filters removed unusually long snoRNAs (e.g., the U3/SNORD3 family; **Figure S2A**) and snoRNAs belonging to large paralogous families as defined by Rfam copy number thresholds^25,26^ (**Figure S2B**). The final network contains 215 box C/D snoRNAs and 166 RBPs, representing ∼65% of expressed human box C/D snoRNAs in snoDB and ∼99% of ENCODE RBPs with high-confidence eCLIP data. Each node was assigned an expression score by binning its highest observed transcript abundance across referenced cell lines into four discrete categories (**Figure S3**).

**Figure 1.**
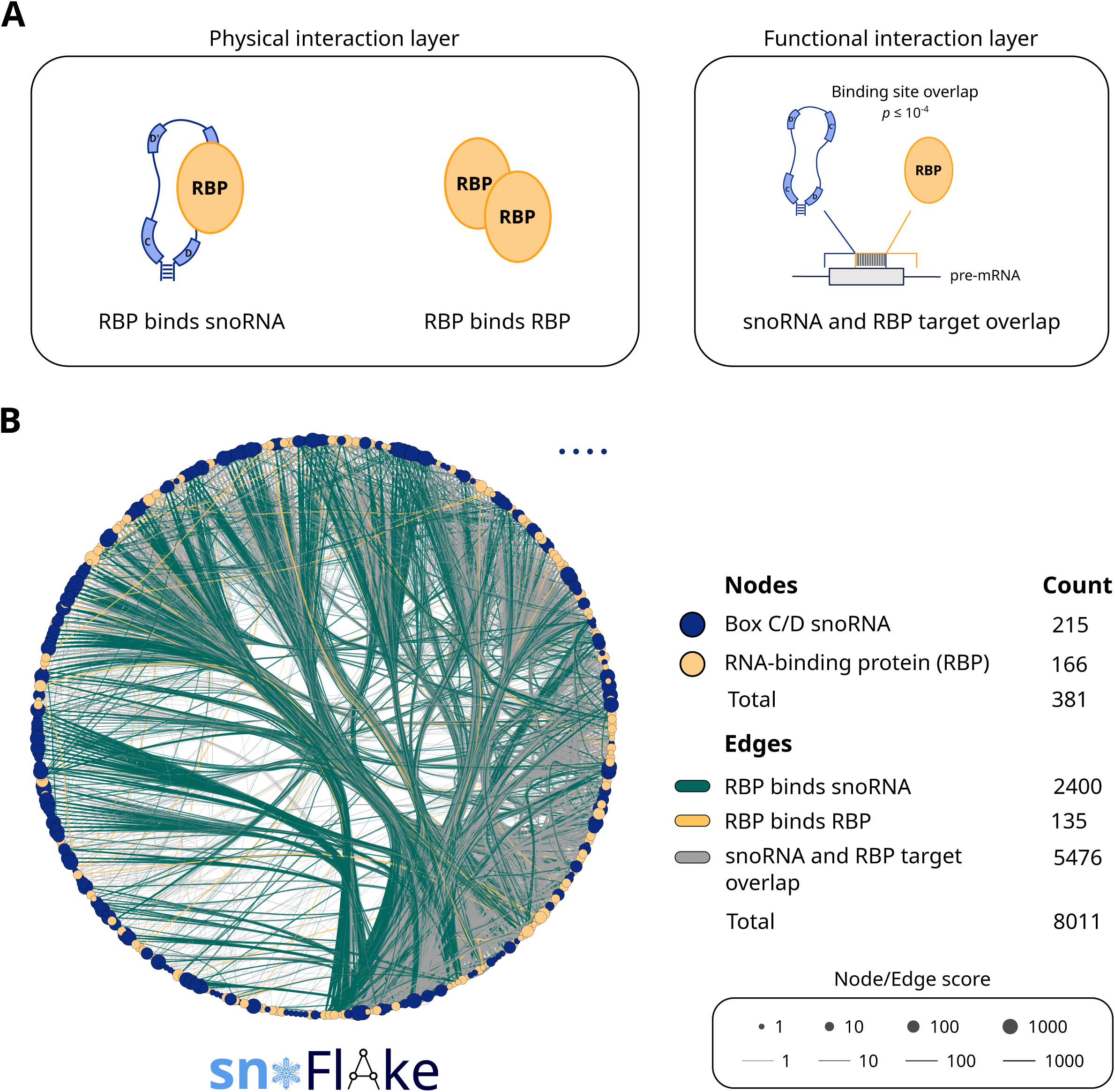
snoFlake: A functional interactome of human box C/D snoRNAs and RNA-binding proteins. **(A)** Schematic overview of the snoFlake network architecture. snoFlake integrates two interaction layers: (1) a physical interaction layer comprising direct RBP–snoRNA binding and RBP–RBP protein interactions and (2) a functional layer defined by co-targeting of shared protein-coding transcripts by both a snoRNA and an RBP. Functional interactions were assigned to snoRNA–RBP pairs exhibiting significant enrichment of overlapping binding sites on shared protein-coding RNA targets (Fisher’s exact test, Benjamini–Hochberg FDR ≤ 0.001, corresponding to *p* ≤ 10⁻⁴; STAR Methods). **(B)** Global view of the snoFlake network visualized using a degree-sorted circular layout in Cytoscape^33^. Nodes represent box C/D snoRNAs (dark purple) and RBPs (orange). Node size reflects the expression score calculated based on the maximum RNA abundance (in TPM) observed across five human cancer cell lines^23,24^ (Figure S3). Larger nodes represent snoRNAs or RBPs with higher RNA abundance. Edges correspond to the three interaction types described in (A) and are displayed in distinct colors. Edge thickness reflects interaction confidence score, with thicker edges indicating stronger statistical support (Figure S4D). See also Figure S1 for the detailed network construction pipeline and Table S1 for snoFlake node and edge annotations.

To capture multiple dimensions of box C/D snoRNA regulation, we modeled three types of relationships (**Figures 1A and S1**). First, we defined physical snoRNA–RBP interactions as binding events identified from ENCODE eCLIP datasets^22^. These interactions mark regions where RBPs directly contact snoRNAs, with the distribution of binding site lengths shown in **Figure S4A**. Second, physical RBP–RBP interactions were derived from the STRING database (v12.0)^27^, representing protein-protein associations that suggest potential co-complex formation. Third, functional snoRNA–RBP interactions were defined as statistically significant overlaps between snoRNA and RBP binding sites on shared protein-coding RNA targets (Fisher’s exact test, *p* ≤ 10^−□^). RBP binding sites were sourced from ENCODE eCLIP^22^ data, while snoRNA targets were compiled from high-throughput RNA–RNA interaction studies (PARIS^28^, PARIS2^10,29^, LIGR-seq^30^, and SPLASH^31^) and supplemented with transcriptome-wide machine-learning predictions from snoGloBe^32^ to mitigate the coverage limitations of experimental approaches. In total, snoFlake incorporates 8,011 edges: 2,400 physical snoRNA–RBP edges, 135 physical RBP–RBP edges, and 5,476 functional snoRNA–RBP edges (**Figure 1B**). Notably, functional interactions outnumber direct binding snoRNA–RBP interactions by 2.3-fold, highlighting the extensive co-targeting relationships between snoRNAs and RBPs. To validate the biological relevance of these functional interactions, we compared snoRNA–RBP pairs with significant target overlap (*n* = 5,476) against non-significant pairs (*n* = 13,358) and found that significant pairs exhibited greater numbers of shared targets and longer overlapping binding regions (Mann–Whitney *U* test, *p* < 0.001; **Figures S4B and S4C**). To enable comparative visualization, each edge was assigned a confidence score by binning interaction significance values into four discrete categories (**Figure S4D**).

Network topology analysis revealed a heterogeneous distribution of interactions across both snoRNAs and RBPs. Among snoRNAs with at least one interaction (*n* = 211), node degrees, defined as the total number of interaction edges connected to each snoRNA, ranged from 1 to 148 (median = 28; **Figure S4E**). RBPs (*n* = 166) showed a similarly wide range of connectivity, spanning 1 to 201 interactions (median = 42; **Figure S4F**). This broad distribution highlights a modular network architecture in which a few highly connected nodes act as potential hubs, while most components participate in fewer interactions. In particular, SNORD23 and SNORD17 displayed the highest degrees among snoRNAs, whereas NOLC1 and AATF were the most highly connected RBPs, suggesting that these molecules may occupy central roles in snoRNA-guided regulatory assemblies.

To our knowledge, no dedicated database currently catalogs snoRNA–RBP interactions, positioning snoFlake as a unique and comprehensive resource for exploring the organization and regulatory potential of snoRNA-protein interactions and their broader impact on post-transcriptional gene regulation. All snoFlake interactions and node-level annotations are provided in **Table S1** and a downloadable interactive network is available as a Cytoscape^33^ session file in **Data S1**.

### Double-edge snoRNA–RBP interactions are enriched for spliceosomal proteins

To prioritize high-confidence and potentially functional snoRNA–RBP relationships, we focused on interactions supported by two independent lines of evidence: physical binding between a snoRNA and an RBP and functional co-targeting of shared protein-coding RNAs. We refer to these combined associations as “double-edge” interactions, represented as parallel edges connecting a snoRNA–RBP pair (**Figure 2A**). In total, we identified 338 double-edge snoRNA–RBP pairs, accounting for ∼14% of all physical snoRNA–RBP interactions and ∼6% of all functional interactions in snoFlake (**Figure 2B**). These pairs involved 95 snoRNAs and 67 RBPs, corresponding to ∼44% of snoRNAs and ∼40% of RBPs in the network (**Figures S5A and S5B**), indicating that double-edge interactions represent a functionally enriched subset of snoRNA–RBP relationships.

**Figure 2.**
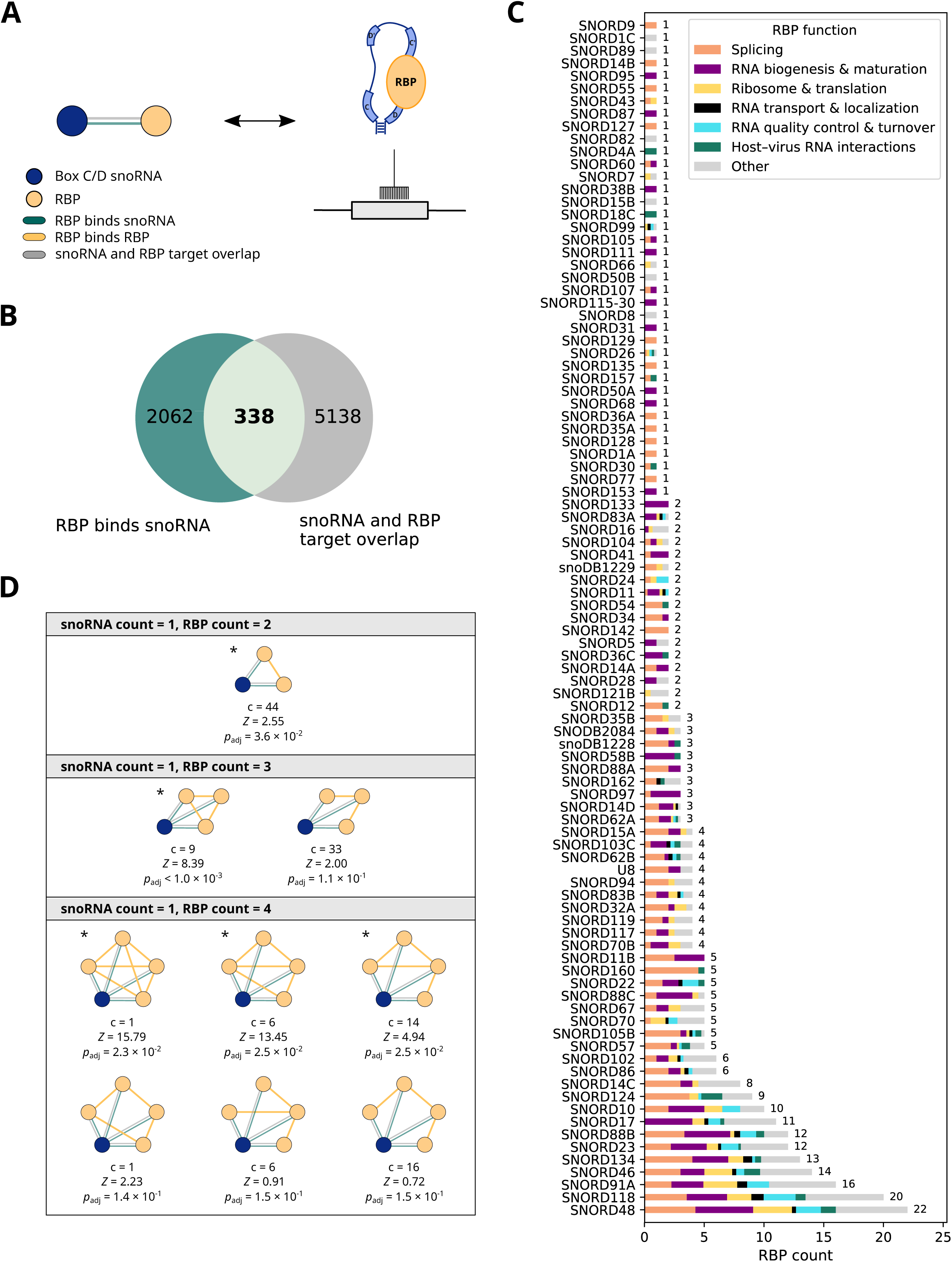
Double-edge motif analysis reveals distinct regulatory snoRNA–RBP complexes. **(A)** Schematic representation of a double-edge interaction in snoFlake, shown as a network diagram (left) and corresponding biological model (right). A double-edge interaction is defined by the presence of both physical binding of an RBP to a box C/D snoRNA (green edge in network diagram) and significant enrichment of overlapping binding sites on shared protein-coding RNA targets (gray edge), suggesting a cooperative regulatory association. **(B)** Venn diagram showing the overlap between physical RBP–snoRNA interactions (green) and snoRNA–RBP target overlap interactions (gray) in snoFlake, with numbers indicating interaction counts. The intersection represents double-edge interactions as defined in (A). **(C)** Bar plot showing the number of RBPs participating in double-edge interactions with each snoRNA (total RBP count indicated to the right of each bar). Bars are segmented and color-coded according to the functional categories represented among the RBPs involved in the corresponding double-edge interactions. For multifunctional RBPs, functional annotations were apportioned equally across categories (1/*n*; *n* is the number of functional categories assigned to that RBP), yielding a proportional breakdown of RBP functions within each bar. Only snoRNAs with at least one double-edge interaction are shown, with each bar representing one snoRNA. **(D)** SnoRNA-centered network motifs present in snoFlake. Motifs were defined as subnetworks in which all snoRNA–RBP pairs are connected by double-edge interactions and at least two RBPs are linked by an RBP–RBP binding edge. Motif enrichment was assessed by comparing observed counts to those in degree-preserving random networks (STAR Methods). Motif counts include all occurrences of a given topology, such that overlapping motifs of lower complexity were counted independently. For each motif type, the observed count (*c*), Z-score (*Z*), and adjusted p-value (*p*_adj_) are shown below each motif. P-values were corrected using the Benjamini–Hochberg method (FDR *<* 0.05). Statistically significant motifs (*p*_adj_ < 0.05) are indicated by an asterisk (*). See also Figure S8 for individual instances of each motif, displayed at the highest-order topology observed for each snoRNA–RBP configuration.

Examining the functional landscape of double-edge interactions, we found that many snoRNAs were associated with multiple RBPs involved in interrelated aspects of RNA metabolism (**Figure 2C**). These included not only orphan snoRNAs but also canonical box C/D snoRNAs such as SNORD48 and SNORD32A, which are typically known for guiding 2 ′ -O-methylation of rRNA^34^. Notably, 58 of the 67 double-edge RBPs (∼87%) interacted with at least one canonical guide snoRNA (**Figure S6**), indicating that these RNAs may commonly participate in regulatory contexts that extend beyond their established roles in RNA modification. Functionally, snoRNAs involved in double-edge interactions showed strong enrichment for RBPs linked to splicing (**Figure 2C**). Approximately 69% (66/95) of these snoRNAs were connected to at least one splicing-annotated RBP, 60% (57/95) to RBPs involved in RNA biogenesis and maturation, and 43% (41/95) to RBPs associated with ribosome biogenesis and translation. These functional categories were not mutually exclusive, reflecting the interconnected nature of RNA metabolic pathways.

Given the prominence of splicing-related interactions, we further examined the 66 snoRNAs that formed double-edge interactions with 22 splicing-associated RBPs (**Figure S5**). Within this group, double-edge interactions were more frequent with core spliceosomal proteins than with splicing regulators or auxiliary factors (96 vs. 24 vs. 7 interactions, respectively)^35–37^. A recurring set of catalytic spliceosomal RBPs dominated this network layer, including U2 snRNP components (SF3B4, SMNDC1)^38,39^ and factors associated with the U5 snRNP and PRP19 complex (PRPF8, EFTUD2, AQR)^40,41^ (**Figures S7A and S7B**), which participate in spliceosome assembly, catalytic activation, and conformational remodeling during the splicing reaction. The prevalence of interactions with these proteins suggests that snoRNAs are not merely passive binding partners or peripheral regulatory factors but may directly interface with the splicing machinery to fine-tune catalytic steps or coordinate RNA-protein remodeling events. These findings align with and extend emerging evidence for snoRNA-guided modulation of alternative splicing^16,18^, pointing to a potentially conserved yet noncanonical functional relationship between snoRNAs and the spliceosome.

### Enriched snoRNA–RBP network motifs reveal modular spliceosomal assemblies

To identify higher-order interaction patterns within double-edge snoRNA–RBP relationships, we focused on snoRNA-centered network motifs, defined as snoRNAs forming double-edge interactions with two or more RBPs that are themselves linked through physical RBP–RBP interactions. Disconnected RBP groups linked to the same snoRNA were treated as separate motifs. Under this definition, the largest motif contained four RBPs. Considering all possible configurations with one snoRNA and two to four RBPs yielded nine distinct motif topologies (**Figure 2D**). Motif enrichment was evaluated by comparing motif counts in snoFlake against those in 1,000 degree-preserving randomized networks (see **STAR Methods**). This conservative null model controls for node connectivity while disrupting higher-order interaction patterns and revealed that five of the nine motif topologies were significantly enriched (*p*_adj_ < 0.05). Enrichment statistics were based on total motif counts across the network, including all occurrences of a given topology and overlapping instances.

In total, we identified 23 distinct motifs, each representing a unique snoRNA–RBP configuration, involving 19 snoRNAs (20% of double-edge snoRNAs) and 18 RBPs (27% of double-edge RBPs; **Figure S8**). Within each motif, RBP partners typically shared functional annotations, suggesting cooperative multiprotein assemblies rather than isolated or incidental pairwise contacts. We therefore classified motifs by biological process based on the dominant function of their RBP components. Splicing represented the dominant category (15/23), followed by ribosome biogenesis and translation (6/23), with the remaining two motifs comprising RBPs of mixed functions. Among the splicing-related motifs, RBPs within the same motif consistently mapped to either the same spliceosomal complex or to complexes known to act in concert during spliceosome assembly.

Specifically, motifs comprised RBPs exclusively from the U2 snRNP, exclusively from the U5 snRNP, or from functionally related pairs such as the U2 and U5 snRNPs^42^ or the U5 snRNP and the PRP19 complex^43^. This co-complex organization within motifs, which was not apparent at the level of individual double-edge interactions, motivated experimental investigation of top-ranked motifs as candidate snoRNA–RBP regulatory complexes.

### snoFlake identifies noncanonical SNORD22 interactions with PRPF8 and EFTUD2

Analysis of the 23 enriched motifs by node and edge scores revealed a top-ranked module centered on SNORD22, an orphan box C/D snoRNA^20^ forming double-edge interactions with two core U5 snRNP proteins, PRPF8 and EFTUD2 (**Figure 3A**). Within this motif, SNORD22 and PRPF8 received high node and edge scores, while EFTUD2 showed intermediate scores, reflecting lower transcript abundance and more limited target overlap with SNORD22.

**Figure 3.**
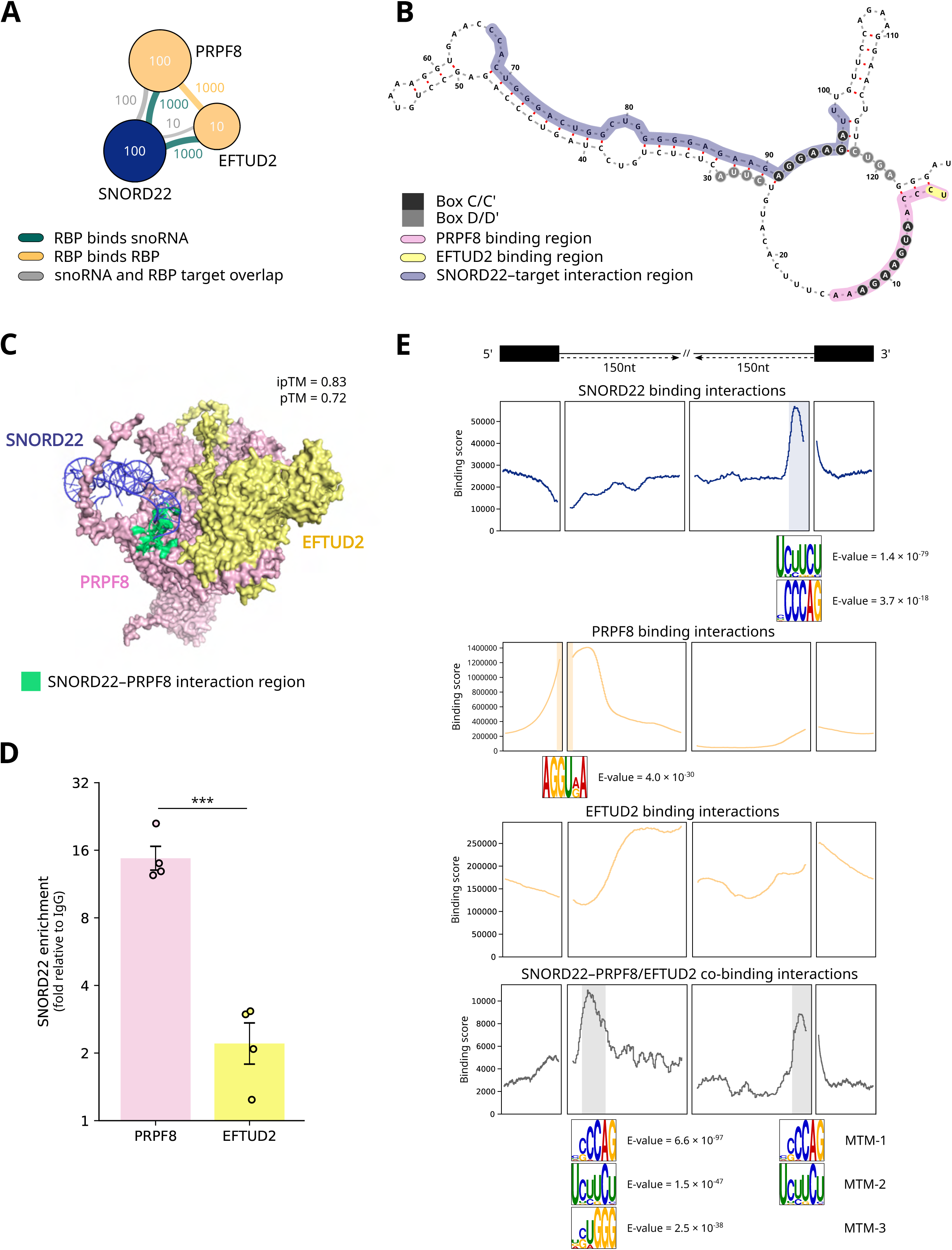
SNORD22 forms a putative complex with U5 snRNP proteins PRPF8 and EFTUD2 and co-binds near splice junctions. **(A)** A statistically significant network motif identified by snoFlake comprising SNORD22 and two U5 snRNP-associated RBPs, PRPF8 and EFTUD2. Node and edge scores are indicated adjacent to the corresponding elements. **(B)** Predicted secondary structure of SNORD22 generated using LinearPartition^72^. PRPF8 and EFTUD2 binding sites on SNORD22, as determined by ENCODE eCLIP data^22^, are highlighted in pink and yellow, respectively. The main predicted target interaction region of SNORD22 with protein-coding RNA targets is indicated in purple. **(C)** AlphaFold3-generated model of the SNORD22–PRPF8–EFTUD2 complex showing the predicted spatial arrangement of the snoRNA and two RBPs. Model confidence is reported as interface (ipTM) and global (pTM) scores^44^. The interface between SNORD22 and PRPF8 is highlighted in green based on residue proximity (< 5 Å) identified using PyMOL^73^. **(D)** Enrichment of SNORD22 detected by RT–qPCR following RNA immunoprecipitation (RIP) in SKOV3.ip1 cells using antibodies against endogenous PRPF8 and EFTUD2. A nonspecific rabbit IgG was used as a negative control. SNORD22 enrichment was calculated relative to the reference RNA RPPH1 and normalized to the IgG control (STAR Methods). Bars represent the geometric mean with back-transformed SEM error bars displayed on a log_2_ scale (*n* = 4 independent biological replicates). Statistical significance was assessed using a two-sided Welch’s *t*-test on log₂-transformed enrichment values (\*\*\**p* < 0.001). **(E)** Transcriptome-wide binding profiles of SNORD22, PRPF8, and EFTUD2. SNORD22 binding interactions were derived from a combination of snoGloBe machine-learning predictions^32^ and high-throughput RNA–RNA interaction datasets^10,28–31^. RBP binding interactions were obtained from ENCODE eCLIP data^22^. Metagene plots show binding distributions across protein-coding transcripts, grouped into four mutually exclusive categories: SNORD22-only, PRPF8-only, EFTUD2-only, and SNORD22 binding sites overlapping with either PRPF8 or EFTUD2. Binding profiles are shown in a ± 150 nt window centered on splice junctions. The y-axis indicates normalized binding score (STAR Methods). Significant sequence motifs were identified using MEME^48^ (E-value ≤ 10^−□^) and are shown below their respective regions of enrichment. In the SNORD22–PRPF8/EFTUD2 co-binding group, enriched motifs are labeled MTM-1, MTM-2, and MTM-3 (MTM: mRNA target motifs) and are referenced in subsequent figures. See also Figure S15 for additional binding profiles of pairwise and triple co-binding combinations of SNORD22, PRPF8, and EFTUD2.

ENCODE eCLIP data^22^ revealed that PRPF8 and EFTUD2 bind the 5′ terminus of SNORD22 (eCLIP *p* < 2.2 × 10⁻¹□), with the PRPF8 footprint spanning nucleotides 1–14 and EFTUD2 binding at positions 1–2 (**Figure 3B**). Chimeric eCLIP data for core box C/D snoRNP proteins (NOP56, NOP58, FBL, and SNU13) showed no significant binding to SNORD22^19^, indicating that SNORD22 likely operates outside the canonical box C/D snoRNP complex or associates only weakly with its core components. Transcriptome-wide mapping showed that the majority (83.8%) of SNORD22 binding interactions involve a 34-nt AG-rich region near box C′, whereas a smaller fraction (14.3%) uses ASE-like regions near boxes D′ or D to bind protein-coding RNAs (**Figures 3B and S9**). This pattern indicates that SNORD22 recognizes its RNA targets through a noncanonical mechanism distinct from canonical box C/D snoRNA base-pairing^4^. AlphaFold3 modeling predicted that PRPF8 forms direct contacts with SNORD22 nucleotides 1–7 (ipTM = 0.83, pTM = 0.72; mean pLDDT = 76.5)^44^, consistent with the eCLIP footprint (**Figure 3C**). To assess the structural plausibility of this interaction within the assembled spliceosome, we superimposed the predicted SNORD22–PRPF8–EFTUD2 complex onto the cryo-EM structure of the U5 snRNP (PDB: 8Q91)^45^. This alignment showed that SNORD22 fits within an accessible U5 snRNP surface with no predicted steric clashes with neighboring components, supporting the structural feasibility of the SNORD22–PRPF8–EFTUD2 interaction within the U5 snRNP particle (**Figure S10**).

To validate the predicted interaction among SNORD22, PRPF8, and EFTUD2, we performed RNA immunoprecipitation (RIP) followed by RT–qPCR. Endogenous PRPF8 and EFTUD2 were immunoprecipitated using specific antibodies and western blot analysis confirmed successful and specific pulldown of each protein, with no detectable signal in the IgG control (**Figures S11A and S11B**). As shown in **Figure 3D**, SNORD22 was significantly enriched in both PRPF8 and EFTUD2 immunoprecipitates, supporting its association with both proteins *in vivo* (fold enrichment over IgG, geometric mean; PRPF8: 14.74; EFTUD2: 2.20; *n* = 4). In fact, SNORD22 enrichment was higher in PRPF8 RIPs, with a ∼6.7-fold greater signal compared to EFTUD2 (Welch’s *t*-test on log - transformed values, *p* = 6.70 × 10^−□^). Together, network motif analysis, structural predictions, and experimental validation consistently identified PRPF8 as the primary SNORD22 binding partner with EFTUD2 in an auxiliary or cooperative role, establishing the SNORD22–PRPF8–EFTUD2 module as a noncanonical snoRNA–spliceosomal assembly associated with the U5 snRNP.

### SNORD22 co-occupies splice sites with U5 snRNP components in a sequence-dependent manner

To investigate whether SNORD22 plays a role in splicing regulation, we analyzed its transcriptome-wide binding profile across protein-coding genes. While the majority of SNORD22 binding sites (85.5%) occurred within introns, positional enrichment analysis revealed a pronounced signal at exonic regions by ∼2.6-fold and at exon-intron junctions by ∼166-fold relative to their expected transcriptomic background (χ*²* test, *p* < 10^−¹□^; **Figure S12**), suggesting preferential targeting of splicing-relevant landmarks. Comparison of SNORD22 binding interactions with those of PRPF8 and EFTUD2 revealed 2,712 sites co-bound by SNORD22 and at least one of the two U5 snRNP proteins (**Figure S13**), corresponding to ∼2% of total binding sites for each factor individually. Genes containing these co-bound regions (*n* = 2,197) were significantly enriched for pathways associated with the cell cycle, stress response, and membrane trafficking (FDR < 0.05; **Figure S14**)^46,47^, indicating that SNORD22–U5 snRNP co-binding marks a specialized layer of post-transcriptional control over specific gene sets.

To refine the spatial context of binding relative to splice junctions, we generated metagene profiles using ±150 nt windows centered on splice sites flanking regions bound by SNORD22, PRPF8, and EFTUD2 (**Figure 3E**). SNORD22-specific binding was enriched at the polypyrimidine tract (PPT) and 3′ splice site (3 ′ SS), representing 21% of its unique interactions. Motif analysis^48^ further identified enrichment for CU-rich PPT-like elements (E-value = 1.4 × 10^−□□^) and canonical 3′SS-like CAG motifs (E-value = 3.7 × 10⁻¹□) at these sites. Consistent with previous spliceosomal studies^40^, PRPF8 exhibited preferential binding near the 5′ splice site (5′SS) in over 90% of unique binding sites, with enrichment for a consensus 5′SS motif (E-value = 4.0 × 10^−30^). EFTUD2 displayed a more diffuse binding pattern (64.2% of interactions shown), reflecting its dynamic repositioning during spliceosome assembly and activation^49^. Aggregated SNORD22 co-binding interactions with PRPF8 and/or EFTUD2 showed a bimodal distribution, with one peak over the 3′SS and PPT and a stronger peak downstream of the 5′SS, collectively accounting for 86.3% of total co-binding interactions. Notably, the majority of co-binding interactions occurred exclusively at either the 5′SS or 3′SS within a given intron, with only ∼2% showing co-occupancy at both splice sites. Surprisingly, both splice site regions were enriched for 3′SS and PPT-like motifs (MTM, mRNA target motif; MTM-1 E-value = 6.6 × 10^−□□^; MTM-2 E-value = 1.5 × 10^−□□^), matching SNORD22’s intrinsic AG-rich target-recognition sequence. Taken together, these findings suggest that SNORD22 may drive co-binding site selection through local sequence context rather than through splice site position. Additional binding profiles and sequence motif analyses for all pairwise and triple combinations not shown in Figure 3E are provided in **Figure S15**.

### SNORD22 promotes inclusion of weakly spliced cassette exons

To evaluate SNORD22’s functional impact on pre-mRNA splicing, we analyzed transcriptome-wide changes following its depletion. Two RNase H-dependent antisense oligonucleotides (ASOs) targeting distinct regions of SNORD22 were transfected into SKOV3.ip1 ovarian cancer cells (RRID: CVCL_0C84)^50^, achieving 40–50% reduction in SNORD22 expression as measured by RT–qPCR (**Figures S16A–C**). Polyadenylated (poly(A)+) RNA-seq libraries were prepared from two biologically independent replicates per ASO and negative control, yielding two knockdown (KD) samples per ASO (four SNORD22 KD samples total) and two control samples for downstream analysis (**Figure 4A**).

**Figure 4.**
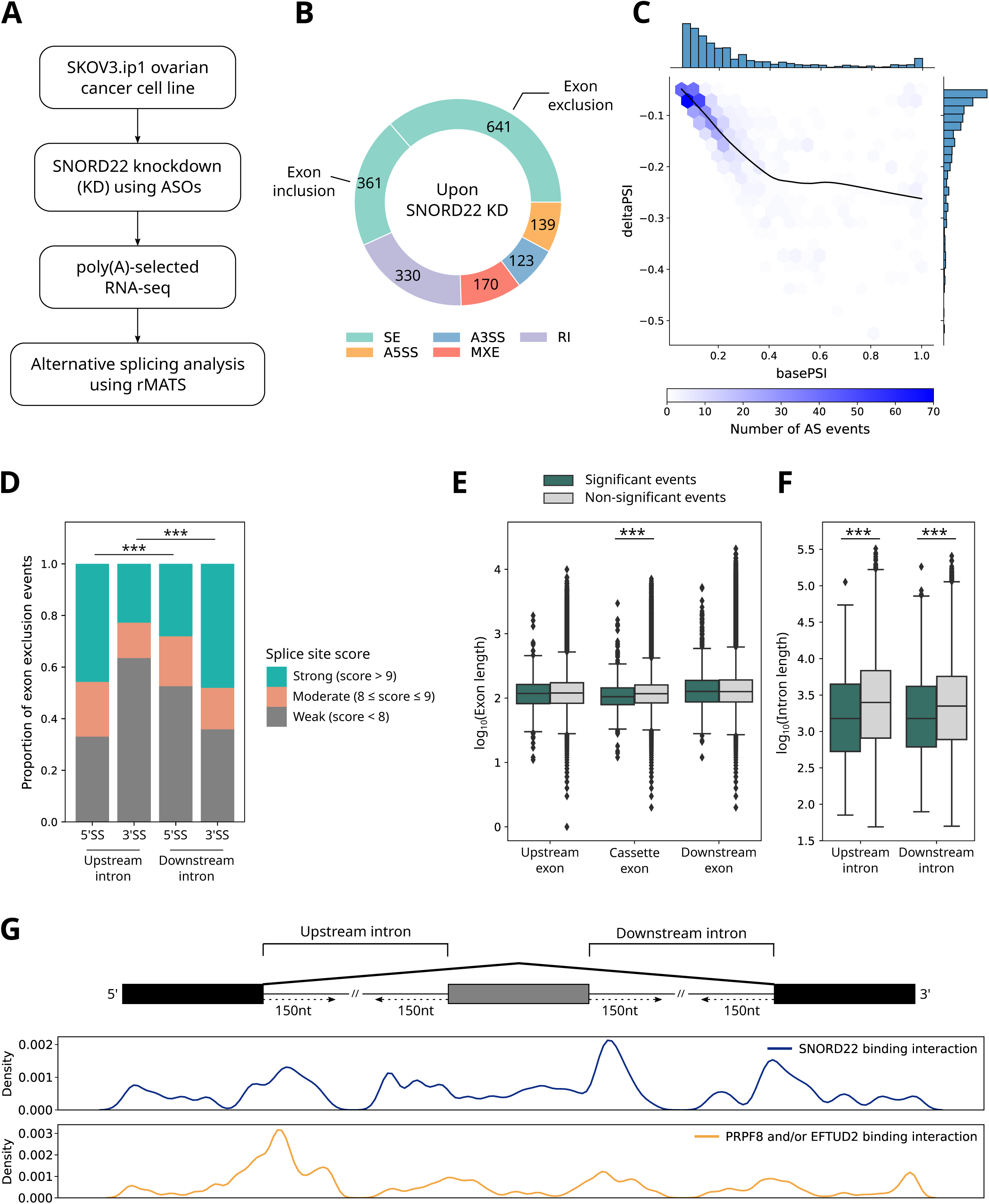
SNORD22 promotes inclusion of suboptimal cassette exons. **(A)** Experimental workflow for the identification of SNORD22-dependent alternative splicing events. SNORD22 was depleted in SKOV3.ip1 ovarian cancer cells using two independent antisense oligonucleotides (ASOs; Figure S16), followed by poly(A)-selected RNA-seq and alternative splicing analysis using rMATS^52^. **(B)** Doughnut plot showing the number of significant alternative splicing events detected by rMATS upon SNORD22 knockdown (KD) (*n* = 1,764). Significant events were defined as those with read count ≥ 10, |ΔPSI| ≥ 0.05, and FDR ≤ 0.01 in genes expressed in at least one condition (KD or control). Events were categorized as skipped exon (SE), alternative 5′ splice site (A5SS), alternative 3′ splice site (A3SS), mutually exclusive exons (MXE), and retained intron (RI), with SE events further categorized as exon inclusion or exon exclusion. **(C)** Hexbin plot showing the relationship between baseline percent spliced in (basePSI; x-axis) and ΔPSI (SNORD22 KD PSI – control PSI; y-axis) for 641 exon exclusion events identified in (B). Each hexagon represents the number of events within that bin, with darker shades indicating higher event density. Marginal histograms show the distributions of basePSI (top) and ΔPSI (right). A locally weighted scatterplot smoothing (LOWESS) curve is overlaid to indicate the global trend. **(D)** Proportional bar plot showing the splice site strength of the 5′ and 3′ splice sites (5′SS and 3′SS) of the upstream and downstream introns flanking the cassette exons excluded upon SNORD22 KD (*n* = 641). Splice site strength was calculated using MaxEntScan^53^ and categorized as weak (score < 8), moderate (8 ≤ score ≤ 9), or strong (score > 9). Pairwise chi-squared tests were performed across the four splice sites with Benjamini–Hochberg correction (\*\*\**p*_adj_ < 0.001). **(E)** Box plots of exon lengths for the 641 exon exclusion events identified in (B). Length distributions are shown for the upstream, cassette, and downstream exons. Events are grouped as significantly alternatively spliced (read count ≥ 10, |ΔPSI| ≥ 0.05 and FDR ≤ 0.01; dark green) or non-significantly spliced (read count ≥ 10, |ΔPSI| < 0.05 or FDR > 0.01; gray). Differences in exon length between groups were assessed separately for each exon type using the Mann–Whitney *U* test with Benjamini–Hochberg correction (\*\*\**p*_adj_ < 0.001). **(F)** As in (E) but showing length distributions for the upstream and downstream introns flanking the cassette exon. **(G)** Binding profiles of SNORD22, PRPF8, and EFTUD2 at SNORD22-sensitive exon exclusion events identified in (B). Binding sites derived from snoGloBe predictions^32^ and HTRRI datasets^10,28–31^ (SNORD22) or ENCODE eCLIP data^22^ (PRPF8 and EFTUD2) are mapped to the upstream exon, cassette exon, downstream exon, and ±150 nt flanking each splice site. Density curves highlight positional enrichment for SNORD22 (purple) and the union of PRPF8 and EFTUD2 binding (orange). Only events with SNORD22 binding within the shown region were included (*n* = 89).

Considering only genes expressed in at least one condition (KD or control), differential expression analysis using DESeq2^51^ identified 811 downregulated and 167 upregulated protein-coding genes in SNORD22 KD samples relative to controls (|log□ fold change| > 1, *p*_adj_ < 0.05; **Figure S17A**). Nevertheless, differentially expressed genes showed no enrichment for SNORD22 co-binding with PRPF8 or EFTUD2 compared to non-differentially expressed genes (**Figure S17B**), indicating that these transcriptional changes most likely occur independently of the SNORD22–U5 snRNP interaction.

To assess the impact of SNORD22 depletion on pre-mRNA splicing, we performed alternative splicing analysis using rMATS^52^, which identified 1,764 significantly altered splicing events (read count ≥ 10, |ΔPSI| ≥ 0.05, FDR ≤ 0.01). The majority of these events (*n* = 1,002) were skipped exons (SE), also known as cassette exons (**Figure 4B**). Among these, 641 showed reduced exon inclusion (hereafter referred to as exon exclusion events) while 361 exhibited increased inclusion. Strikingly, exclusion events were concentrated among cassette exons with low basal Percent Spliced In (PSI) values, where approximately 66% (425/641) had PSI < 0.25 under control conditions (median PSI = 0.178; **Figure 4C**). Consistent with this weak basal inclusion level, ΔPSI values were modest (typically between –0.05 and –0.15) due to a floor effect. These findings suggest that SNORD22 may promote inclusion of weakly spliced cassette exons by enhancing their recognition by the spliceosome.

To explore the structural features shared by these 641 SNORD22-sensitive cassette exons, we used MaxEntScan^53^ to score the strength of the four splice sites flanking each cassette exon—the upstream intron 5′SS and 3′SS and the downstream intron 5′SS and 3′SS—and classified them as weak (score < 8), moderate (8 ≤ score ≤ 9), or strong (score > 9). The cassette exon-proximal splice sites (upstream intron 3 ′ SS and downstream intron 5 ′ SS) had a significantly higher proportion of weak scores compared to their distal counterparts (downstream intron 3′SS and upstream intron 5′SS, respectively; χ² test, *p*_adj_ < 0.001; **Figure 4D**). In total, 83% (534/641) of cassette exons had at least one weak proximal splice site: 63% (407/641) had a weak upstream intron 3 ′SS, 53% (337/641) a weak downstream intron 5′SS, and 33% (210/641) had both (categories not mutually exclusive). Compared to non-differentially spliced events upon SNORD22 KD, these exons were also shorter (median 105 nt vs. 117 nt, Mann–Whitney *U* test, *p*_adj_ < 0.001; **Figure 4E**), exhibited lower GC content (median 0.474 vs. 0.489, *p*_adj_ < 0.01; **Figure S18**), and were flanked by shorter introns (upstream intron median 1,506 nt vs. 2,498 nt; downstream intron median 1,507 nt vs. 2,233 nt; *p*_adj_ < 0.001; **Figure 4F**). Collectively, these suboptimal structural features likely contribute to inefficient exon recognition by the spliceosome^54,55^, explaining their characteristically low PSI under basal conditions.

We next investigated whether these SNORD22-sensitive cassette exons exhibit specific co-binding patterns between SNORD22 and the U5 snRNP components PRPF8 and EFTUD2. In line with our earlier observation that SNORD22–PRPF8/EFTUD2 co-binding occurs preferentially near splice sites (**Figure 3E**), global co-binding across the entire splicing event region was not significantly enriched among exon exclusion events compared with non-differentially spliced events following SNORD22 KD (Fisher’s exact test, *p* = 0.097; **Figure S19**). We therefore focused on events with SNORD22 binding near splice sites, which are most likely to reflect direct splice-site-level regulation. Of the 641 exon exclusion events, 274 (43%) showed SNORD22 binding within the splicing event region, of which 89 (32%) had SNORD22 binding in the exon or within 150 nt of a splice site. Using these 89 events, we generated metagene binding profiles of SNORD22, PRPF8, and EFTUD2 across ±150 nt windows centered on the splice sites of the cassette exon and its flanking exons (**Figure 4G**). SNORD22 binding was detected at splice sites of both flanking introns, but was preferentially enriched in the downstream intron, with 1.6-fold and 1.4-fold enrichment at the downstream 5 ′ SS and 3′ SS, respectively. In contrast, PRPF8 and EFTUD2 occupancy was 2.6-fold lower at the downstream 5′SS than at the upstream site, indicating reduced U5 snRNP engagement at this junction. Binding at the 3′SS was low for both introns. These asymmetric binding patterns suggest that SNORD22 is preferentially enriched at the downstream intron in regions where U5 snRNP occupancy is reduced. Taken together, these findings support a model in which SNORD22 stabilizes U5 snRNP association at weak cassette exons, thereby facilitating exon inclusion under conditions where basal U5 recruitment is limited.

### Representative loci reveal functional diversity of SNORD22-dependent exon inclusion

To illustrate the functional consequences of SNORD22-dependent regulation of suboptimal cassette exons, we examined two representative loci. *SRRT* (also known as *ARS2*) encodes a cofactor of the nuclear cap-binding complex with essential roles in RNA processing and miRNA biogenesis^56^. Differential splicing analysis^52^ identified a cassette exon in *SRRT* whose inclusion was significantly reduced upon SNORD22 KD (mean control PSI = 0.191, mean KD PSI = 0.075; ΔPSI = −0.117; FDR = 5.9 × 10^−11^; **Figures 5A and S20A**). This reduction in exon inclusion was independently confirmed by ddPCR (**Figure S20B**). The SNORD22-sensitive cassette exon in *SRRT* is flanked by a weak upstream 3′SS (MaxEntScan score 3.88; **Figure S20C**)^53^, which likely contributes to inefficient spliceosomal recognition and its characteristically low basal PSI. SNORD22, PRPF8, and EFTUD2 co-bind two regions within the intron immediately downstream of the exon, one near the 5′SS and a second further downstream in the same intron. At both positions, SNORD22 binds the MTM-2 motif through its AG-rich target-recognition sequence (**Figure S20D**). Inclusion of this exon is predicted to introduce a premature termination codon (PTC) more than 50 nt upstream of the final exon-exon junction^57,58^, producing an NMD-sensitive isoform markedly shorter than the canonical transcript. Despite its low basal PSI, this exon is highly conserved across vertebrates (mean PhastCons100way score 0.98)^59,60^, suggesting strong evolutionary constraint. These features highlight *SRRT* as a representative example in which SNORD22 co-binds with U5 snRNP components near downstream intron splice sites to promote inclusion of a weak cassette exon, potentially directing a subset of transcripts toward NMD.

**Figure 5.**
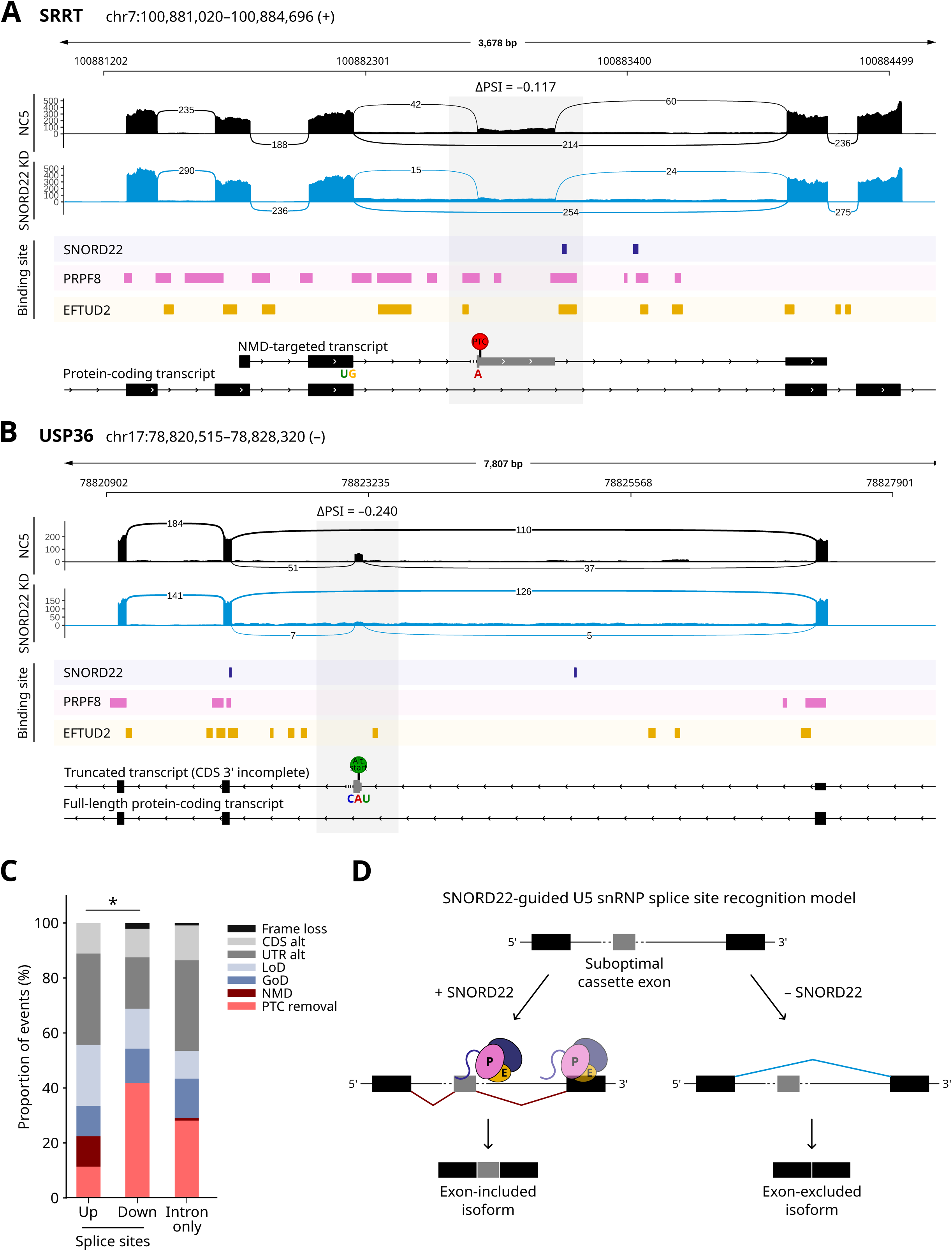
Representative loci reveal functional diversity of SNORD22-dependent exon inclusion. **(A)** Sashimi plot of the *SRRT* locus showing a cassette exon that is preferentially excluded upon SNORD22 knockdown (KD). Read coverage (y-axis) represents the mean across two negative control (NC5; black) and four SNORD22 KD (blue) samples, respectively. Numbers on splice junction arcs indicate the average number of reads supporting each junction. The cassette exon is highlighted with a gray rectangle. Binding profiles for SNORD22, PRPF8, and EFTUD2 are shown below, aligned with *SRRT* transcript annotations. Inclusion of the cassette exon introduces a premature termination codon (PTC), generating a transcript predicted to be targeted by nonsense-mediated decay (NMD). The exon-excluded form encodes the canonical *SRRT* protein-coding transcript. See also Figure S20 for more details. **(B)** Sashimi plot of the *USP36* locus as in (A). Inclusion of the cassette exon results in usage of an alternative start codon, producing a truncated transcript annotated as CDS 3′ incomplete^62^. Skipping of the exon preserves the full-length protein-coding *USP36* transcript. See also Figure S21 for more details. **(C)** Proportion of predicted functional consequences of exon exclusion events following SNORD22 KD, determined using SpliceDecoder^74^. Only splicing events bound by SNORD22 were included. Events were grouped by SNORD22 binding location: upstream intron splice sites (Up; n = 18), downstream intron splice sites (Down; n = 48), or intronic regions only (Intron only; n = 118). Predicted functional consequences were classified as PTC removal, NMD, gain of domain (GoD), loss of domain (LoD), altered UTR (UTR alt), altered CDS (CDS alt), or frame loss. Statistical significance of the difference in PTC removal frequency between upstream and downstream intron splice site groups was assessed using a two-sided Fisher’s exact test (\**p* < 0.05). **(D)** Proposed model of SNORD22-facilitated U5 snRNP splice site recognition. In the presence of SNORD22 (curved line bound to PRPF8), PRPF8 (pink oval) and EFTUD2 (orange oval) are recruited to the downstream intron splice sites of suboptimal cassette exons, promoting exon inclusion. The dark purple oval represents other U5 snRNP components. In the absence of SNORD22, U5 snRNP recruitment is impaired, resulting in exon skipping.

*USP36* encodes a nucleolar deubiquitinase that acts on FBL and other ribosome biogenesis factors^61^, providing a potential link between splicing regulation and box C/D snoRNP biology. SNORD22 KD promoted exclusion of a weakly included cassette exon (mean control PSI = 0.285, mean KD PSI = 0.045; ΔPSI = −0.240; FDR < 2.2 × 10^−16^; **Figures 5B and S21A**), which was independently confirmed by ddPCR (**Figure S21B**). This exon is 72 nt long and has relatively low GC content (GC = 45.8%), consistent with features shared by SNORD22-regulated cassette exons. MaxEntScan scores indicated weak 5′SS in both the upstream and downstream introns flanking the cassette exon (upstream 5′SS: 6.68, downstream 5′SS: 6.41; **Figure S21C**)^53^, contributing to the suboptimal splicing environment of the exon. Binding profiles of SNORD22, PRPF8, and EFTUD2 revealed co-localization near the downstream intron 3 ′ SS. At this position, SNORD22 was predicted to base-pair with MTM-1 and MTM-2 motifs via the 5′ portion of its target-recognition sequence, with additional contacts contributed by adjacent upstream nucleotides (**Figure S21D**). Inclusion of this exon is annotated as introducing an alternative start codon and altering the N-terminus of the predicted coding sequence such that the USP catalytic domain begins further downstream than in the canonical isoform (**Figure S21E**)^62^. However, the corresponding Ensembl transcript is classified as CDS 3 ′ incomplete within a truncated transcript, raising the possibility that this isoform may be nonproductive. *USP36* thus illustrates how SNORD22-dependent inclusion of a suboptimal cassette exon can remodel transcript architecture with consequences for coding potential.

Collectively, these two loci exemplify a recurring regulatory configuration in which SNORD22 co-binds with PRPF8 and EFTUD2 near downstream intron splice sites of suboptimal cassette exons, promoting their inclusion and altering transcript stability and coding potential. To determine whether these effects extend beyond individual examples, we analyzed the predicted consequences of exon exclusion events previously identified as SNORD22-bound within the splicing event region (**Figure 5C**). These events produced diverse transcript outcomes, including removal of premature termination codons (PTCs), protein domain alterations, and changes in coding sequences and UTR structure.

Interestingly, events with SNORD22 binding at downstream intron splice sites showed a higher frequency of PTC removal outcomes (42%) compared with events with upstream splice site binding (11%) (Fisher’s exact test, *p* = 0.021). This enrichment is surprising given that previous transcriptome-wide analyses indicate that AS-NMD isoforms occur in ∼12% of human RefSeq genes^63^. Overall, these observations support a model in which SNORD22 acts as a sequence-specific auxiliary factor that reinforces U5 snRNP engagement at downstream intron splice sites in suboptimal splicing contexts, enabling inclusion of otherwise poorly recognized cassette exons and thereby reshaping transcript stability and coding potential (**Figure 5D**).

## Discussion

SnoRNAs have traditionally been viewed as stable components of canonical snoRNP complexes dedicated to site-specific rRNA and snRNA modification. Here, we challenge this view by showing that individual box C/D snoRNAs can assemble into distinct regulatory complexes with diverse RBPs, thereby extending their functional repertoire into post-transcriptional regulation. To systematically define these assemblies, we developed snoFlake, a human box C/D snoRNA-centric interaction network that integrates physical interactions (snoRNA–RBP and RBP–RBP) with functional snoRNA–RBP co-targeting associations (**Figure 1**). Although several RBP–RNA interaction databases exist (e.g., POSTAR3^64^, starBase^65^, and RNAInter^66^), they provide limited support for snoRNA-focused analyses. snoFlake addresses this gap by integrating multiple layers of interaction evidence within a unified framework centered on box C/D snoRNAs. A key feature of snoFlake is the definition of double-edge interactions, which require both direct snoRNA–RBP binding and co-targeting of shared RNA substrates (**Figure 2A**). This dual-evidence model prioritizes snoRNA–RBP pairs most likely to participate in cooperative regulatory assemblies and enables the detection of higher-order interaction patterns that would not be apparent from individual datasets alone. Motif enrichment analysis identified 23 snoRNA-centered network motifs linking snoRNAs with RBPs involved in diverse aspects of RNA biology, including ribosome biogenesis, splicing, and RNA maturation (**Figure S8**). The strong enrichment of spliceosomal RBPs within these motifs indicates that the organization of spliceosome-associated regulatory assemblies is a prominent and recurrent feature of the box C/D snoRNA interactome.

As a proof-of-principle, we characterized the top-ranked SNORD22–PRPF8–EFTUD2 module, illustrating how snoFlake-derived motifs can reveal previously unrecognized snoRNA–RBP regulatory complexes. Our findings expand the conceptual framework of snoRNA function in two key directions. First, we show that a box C/D snoRNA can act as a sequence-specific regulator of pre-mRNA splicing. Unlike SNORD115^18^, which uses canonical antisense elements to base-pair with target RNAs, SNORD22 recognizes pyrimidine-rich motifs near splice junctions through an AG-rich segment overlapping its box C′ (**Figures 3B and 3E**). This noncanonical targeting mechanism appears to be driven primarily by local sequence context rather than splice site position, consistent with the enrichment of CU-rich sequences flanking both 5′ and 3′ splice sites. Second, our results suggest that a snoRNA can reinforce spliceosomal engagement at a subset of suboptimal cassette exons. SNORD22, PRPF8, and EFTUD2 show spatially asymmetric binding patterns, with SNORD22 preferentially enriched in the downstream intron of cassette exons at sites where PRPF8 and EFTUD2 occupancy is reduced (**Figure 4G**). This architecture suggests that SNORD22 stabilizes U5 snRNP engagement at splice sites with limited recognition potential, thereby promoting inclusion of structurally fragile exons.

The *SRRT* and *USP36* loci illustrate the functional consequences of SNORD22-dependent exon inclusion at individual genes (**Figures 5A and 5B**). In *SRRT*, SNORD22 promotes inclusion of a highly conserved poison exon, generating NMD-sensitive isoforms that likely contribute to mRNA quality control through AS-NMD coupling. In *USP36*, exon inclusion produces a transcript with an altered N-terminus predicted to compromise the USP catalytic domain. Given *USP36*’s role in stabilizing FBL and other nucleolar factors^61^, this observation raises the possibility of a feedback mechanism linking SNORD22-mediated splicing to nucleolar homeostasis and box C/D snoRNP biogenesis, although further experimental validation will be required. More broadly, analysis of SNORD22-regulated cassette exons revealed diverse functional consequences of exon inclusion, including NMD-sensitive transcripts, altered coding sequences, and changes in untranslated regions. These observations indicate that SNORD22 primarily acts at the level of splice site engagement rather than dictating a specific transcript outcome. Instead, the functional consequences of exon inclusion depend on the structural and coding properties of the exon itself.

This SNORD22-mediated stabilization of U5 snRNP engagement at weak cassette exons complements and extends prior observations of snoRNA involvement in splicing. For example, SNORD88B has been proposed to modulate alternative splicing through recruitment of splicing factors^67^, although the mechanism remains poorly defined. In contrast, the SNORD22–PRPF8–EFTUD2 module provides a more fully resolved example supported by computationally predicted and experimentally validated interaction data, spatially patterned binding near splice sites, and transcriptome-wide splicing changes. Notably, most known splicing regulators influence early spliceosome assembly by modulating splice site recognition by U1 or U2 snRNPs^68,69^. In contrast, the SNORD22 module appears to act at the level of U5 snRNP engagement, suggesting that snoRNAs may influence later stages of spliceosome assembly associated with exon alignment and catalytic activation^43,45^.

The strong conservation of SNORD22 across vertebrates (mean PhastCons100way score 0.86)^59^ and its consistently high expression across diverse human tissues and cell lines (mean expression 694 TPM)^20^ suggest that this mechanism reflects an evolutionarily maintained regulatory function. Beyond SNORD22, snoFlake identifies SNORD12 and SNORD105B as additional candidates forming motifs with PRPF8 and EFTUD2 (**Figure S8**), indicating that snoRNA-guided modulation of U5 snRNP engagement may extend to other members of the box C/D snoRNA family.

More broadly, snoFlake highlights the functional versatility of box C/D snoRNAs through their participation in multiple, distinct RBP modules. For example, SNORD46 and SNORD134 form one module with ribosome biogenesis factors and another with U2 snRNP components, suggesting dual roles in ribosome assembly and splicing. This modular organization may enable snoRNAs to dynamically reconfigure their protein partners and RNA targets in response to developmental cues, stress conditions, or cell-type-specific requirements. Several cancer-associated snoRNAs, including SNORD78 and SNORD76^70,71^, are also present in snoFlake and can now be connected to specific snoRNA–RBP modules and transcript targets, providing a framework for understanding the regulatory mechanisms of disease-associated snoRNAs.

### Limitations of the study

While snoFlake provides insights into the noncanonical functional landscape of box C/D snoRNAs, several limitations should be considered. First, snoFlake depends on the scope and composition of currently available datasets and is therefore biased toward splicing-related RBPs as the ENCODE RBP collection^22^ is enriched for this class of proteins. In addition, its coverage is limited to a relatively small number of cell lines and experimental conditions, which may overlook snoRNA–RBP complexes specific to particular cell types or physiological contexts. Furthermore, in the absence of a gold-standard reference set of snoRNA–RBP assemblies, network predictions currently rely on biological plausibility and targeted experimental validation, as illustrated here for SNORD22. Nevertheless, snoFlake is designed to be iteratively refined as new datasets become available, including cell-type-specific eCLIP data, RNA–RNA interactomes, and systematic RNP reconstitution studies.

## Conclusion

By integrating physical and functional interaction data, snoFlake uncovers a modular architecture of snoRNA-centered regulatory assemblies and provides a systematic framework for identifying noncanonical snoRNA functions. Using SNORD22 as a proof-of-principle, we show that individual box C/D snoRNAs can act as sequence-specific scaffolds that reinforce spliceosomal engagement at suboptimal splice sites and thereby influence splicing outcomes and transcript fate. These findings highlight snoRNAs as previously underappreciated organizers of RNA-protein regulatory assemblies and suggest that snoRNA-guided modulation of spliceosomal activity may represent a broader regulatory mechanism in RNA biology. As additional interaction datasets become available, snoFlake provides a scalable framework for systematically uncovering snoRNA-guided regulatory modules across diverse RNA metabolic pathways.

## Resource availability

### Lead contact

Requests for further information and resources should be directed to and will be fulfilled by the lead contact, Michelle S. Scott.

### Materials availability

This study did not generate new unique reagents.

### Data and code availability

- This paper analyzes existing, publicly available data, which are listed in the key resources table.
- All snoFlake interaction data and node-level annotations are provided in **Table S1** and an interactive Cytoscape session file containing the full network is available as **Data S1**.
- SNORD22 knockdown poly(A)-selected RNA-seq data generated in SKOV3.ip1 cells have been deposited at GEO: GSE322747 and are publicly available as of the date of publication.
- The snoFlake construction pipeline is available as a Snakemake workflow at https://github.com/scottgroup/snoFlake (including the custom annotation file Homo_sapiens.GRCh38.110_snoRNAs_tRNAs.sorted.gtf).
- Snakemake workflows for TGIRT-seq, poly(A)-selected RNA-seq, and HTRRI analyses are available at https://github.com/kristinassong (repositories: TGIRTseq, RNAseq, and DuplexDiscoverer-Snakemake).
- Any additional information required to reanalyze the data reported in this paper is available from the lead contact upon request.

## Acknowledgements

This work was supported by a CIHR grant (PJT 479838) to M.S.S. and S.A.E. and a team FRQ-NT grant to M.S.S. and S.A.E. K.S.S. was supported by FRQ-NT Master’s and NSERC Doctoral scholarships. L.F.G. was supported by an FRQ-S Doctoral scholarship. M.S.S. holds a Tier 1 Canada Research Chair in Bioinformatics of Noncoding RNA. The authors would like to thank members of the Scott and Abou Elela groups for helpful discussions and the Digital Research Alliance of Canada for providing state-of-the-art computing infrastructure.

## Author contributions

M.S.S. and S.A.E. conceived the study. K.S.S., M.S.S., and S.A.E. designed the experiments. K.S.S. assembled and integrated the datasets with the help of G.D.F., built the network, and analyzed the SNORD22 KD sequencing datasets. M.C., L.F.G., B.Y., and V.K.S. performed the experimental validation. L.F.G. prepared the SNORD22 KD samples. K.S.S. interpreted the data with the help of M.S.S. M.S.S. and K.S.S. wrote the manuscript, and all other authors provided feedback on the manuscript. All authors read and approved the final manuscript.

## STAR Methods

### Experimental model and study participant details

#### Cell lines

The ovarian serous cystadenocarcinoma cell line SKOV3.ip1 (RRID: CVCL_0C84)^50^ was cultured in DMEM/F12 (50/50) medium supplemented with 10% (v/v) fetal bovine serum (FBS) and 2 mM L-glutamine. Cell passaging was performed as recommended by the American Type Culture Collection (ATCC) and no more than 20 passages were used. Cells were periodically tested for mycoplasma contamination by the RNomics Platform at the Université de Sherbrooke. Cell cultures were maintained at 37°C under 5% CO_2_.

### Method details

#### snoFlake network construction and analysis

##### Data curation

The list of human box C/D snoRNAs was collected from snoDB (v2.0)^20^ and filtered based on RNA abundance levels across five TGIRT-seq cancer cell lines (HCT116, MCF7, PC3, TOV112D, SKOV3.ip1) from previous studies^23,24^. Analysis of these TGIRT-seq datasets is described in the following subsections. To exclude unusually long snoRNAs and highly multi-copy families that could confound network and motif analyses, only expressed box C/D snoRNAs ≤ 250 nt in length with ≤ 6 expressed copies were retained in snoFlake (**Figure S2**). The number of expressed copies was determined using Rfam (v15.0)^26^ by identifying all human snoRNAs belonging to the same Rfam family and counting those that met the expression threshold. Human RBPs were obtained from the ENCODE project for those with eCLIP assays in K562 and HepG2 cell lines^22^ and filtered based on mRNA abundance as a proxy for protein availability.

SnoRNA–RNA interactions were compiled from both high-throughput experimental and computational datasets. Analysis of high-throughput RNA–RNA interaction (HTRRI) datasets is described in the following subsections. Computationally predicted interactions were obtained through snoGloBe, a gradient-boosting box C/D snoRNA–RNA interaction predictor^32^. snoGloBe was applied to identify interactions between all box C/D snoRNAs in snoFlake and all protein-coding transcripts (Ensembl biotype: protein_coding) using the custom annotation file Homo_sapiens.GRCh38.110_snoRNAs_tRNAs.sorted.gtf (Ensembl GRCh38 release 110^62^ with additional snoRNA and tRNA annotations; available at https://github.com/scottgroup/snoFlake), with the following parameters: -s 2 -c 25000000 -t 0.95 --seq -m -w 3. HTRRI and snoGloBe-predicted interactions were merged using bedtools merge -s through pybedtools (v0.9.1)^75^ to create a comprehensive interaction dataset for snoFlake.

RBP–RNA interactions were extracted from ENCODE eCLIP datasets^22^ in both K562 and HepG2 cell lines and filtered for significance based on p-value and fold enrichment thresholds following ENCODE eCLIP guidelines. Interactions were first merged across two biological replicates within each cell line, then merged between K562 and HepG2 cell lines using bedtools merge -s (v2.31.1)^76^, resulting in a single unified binding interaction dataset per RBP. RBP–RBP interactions were obtained from STRING (v12.0)^27^, specifically filtering for physical links at high confidence.

Unless otherwise stated, all expression-based filtering thresholds and statistical cutoffs applied during data curation are described in the Quantification and Statistical Analysis section.

Known modification sites on rRNA and snRNA targeted by box C/D snoRNAs were obtained from snoRNA-LBME-db^77^ and snoDB (v2.0)^20^. Functional annotations for RBPs were initially obtained from ENCORE (v2.2.0.6)^22^. RBPs with incomplete annotations were supplemented using GO Biological Process and GO Cellular Component terms from UniProt^35^. Functional categories were then manually consolidated into broader groups (e.g., splicing, RNA biogenesis and maturation, RNA transport and localization) to reduce redundancy. Splicing-related annotations were further refined using the Spliceosome Database^36^, UniProt and a previous study on the transcriptome-wide splicing network^37^. These consolidated annotations were used to characterize the functional composition of RBPs participating in network motifs.

##### TGIRT-seq processing pipeline

To simultaneously quantify snoRNAs and protein-coding transcripts, TGIRT-seq data from five human cancer cell lines (HCT116, MCF7, PC3, TOV112D, SKOV3.ip1)^23,24^ were reanalyzed using a Snakemake workflow available at https://github.com/kristinassong/TGIRTseq. TGIRT-seq uses bacterial thermostable group II intron reverse transcriptases for library preparation, enabling accurate quantification of highly structured RNAs such as tRNAs and snoRNAs^23,78^. Paired-end TGIRT-seq reads were trimmed with Trimmomatic (ILLUMINACLIP:resources/Adapters-PE_NextSeq.fa:2:12:10:8:true TRAILING:30 LEADING:30 MINLEN:20; v0.40)^79^ and raw and trimmed read quality was assessed with FastQC (v0.12.1)^80^. Reads were aligned with STAR (v2.7.11b)^81^ to a CoCo-corrected custom annotation file described above (Homo_sapiens.GRCh38.110_snoRNAs_tRNAs.sorted.gtf), which accounts for nested and multi-mapped genes to enable improved quantification of snoRNAs^62,82^. STAR was run with the following options: --outReadsUnmapped Fastx --outFilterType BySJout --outStd Log --outSAMunmapped None --outSAMtype BAM SortedByCoordinate --outFilterScoreMinOverLread 0.3 --outFilterMatchNminOverLread 0.3 --outFilterMultimapNmax 100 --winAnchorMultimapNmax 100 --alignEndsProtrude 5 ConcordantPair. Abundance levels for snoRNAs and protein-coding RNAs were then estimated as transcripts per million (TPM) using CoCo correct count (cc) with --countType both --strand 1 --paired.

##### High-throughput RNA–RNA interaction analysis

HTRRI datasets generated using PARIS^28^, PARIS2^10,29^, LIGR-seq^30^, and SPLASH^31^ methodologies across diverse human cell types were obtained from the NCBI Sequence Read Archive (SRA) using SRA Explorer^83^ (accessions: PARIS, SRR2814761-SRR2814765; PARIS2, SRR11624581-SRR11624589, SRR11951629, SRR24883901-SRR24883909; LIGR-seq, SRR3361013 and SRR3361017; SPLASH, SRR3404924-SRR3404928 and SRR3404936-SRR3404943).

3 ′ adapters were trimmed with Trimmomatic (v0.40)^79^ using the parameters ILLUMINACLIP:resources/adapters.fa:3:20:10 SLIDINGWINDOW:4:20 MINLEN:18. PCR duplicates were removed with readCollapse.pl from the icSHAPE pipeline^84^. Reads were further trimmed for 5 ′ adapters using Trimmomatic with HEADCROP:17 MINLEN:20. Read quality for raw and processed libraries was assessed with FastQC (v0.12.1)^80^. All processed reads were aligned to the CoCo-corrected annotation described above using STAR (v2.7.11b)^81,82^. STAR indices were generated with --genomeSAindexNbases 14 --genomeChrBinNbits 18. Alignments were run with the following options: --genomeLoad NoSharedMemory --outReadsUnmapped Fastx --outFilterMultimapNmax 10 --outFilterScoreMinOverLread 0 --outSAMattributes All --outSAMtype BAM Unsorted SortedByCoordinate --alignIntronMin 1 –scoreGap 0 --scoreGapNoncan 0 --scoreGapGCAG 0 --scoreGapATAC 0 --scoreGenomicLengthLog2scale −1 --chimOutType Junctions WithinBAM HardClip --chimSegmentMin 5 --chimJunctionOverhangMin 5 --chimScoreJunctionNonGTAG 0 --chimScoreDropMax 80 --chimNonchimScoreDropMin 20 --chimMultimapNmax 5 –limitOutSJcollapsed 10000000 --limitIObufferSize 1500000000 --quantMode GeneCounts.

Processed BAM files were analyzed using DuplexDiscoverer^85^ with default parameters to identify RNA–RNA duplexes. Duplexes detected in PARIS, PARIS2, LIGR-seq, and SPLASH datasets were aggregated so that the support for a given interaction corresponds to the total number of supporting chimeric reads across all samples. The complete Snakemake workflow^21^ for HTRRI processing is available at https://github.com/kristinassong/DuplexDiscoverer-Snakemake.

##### Network construction

snoFlake was built from all human box C/D snoRNAs and RBPs that passed the expression filter described above, represented as two distinct node types. “RBP binds snoRNA” edges were defined from ENCODE eCLIP RBP–RNA interactions^22^ by retaining binding sites that overlapped snoRNA coordinates by at least 1 nt. “RBP binds RBP” edges were taken directly from STRING (v12.0)^27^, restricted to RBPs present in the network. Self-interactions were excluded. Computation of “snoRNA and RBP target overlap” edges is described in the following subsection. In total, snoFlake comprises two node types and three edge types, which are illustrated in different colors (**Figure 1B**).

Node scores were based on the maximum RNA abundance of each snoRNA or RBP across the five cancer cell lines used for initial expression filtering. Edge scores reflect interaction confidence and were derived as follows: ENCODE eCLIP −log_10_(*p*)^22^ for “RBP binds snoRNA” edges, STRING combined score^27^ for “RBP binds RBP” edges, and −log_10_(*p*) values from the overlap analysis described below for “snoRNA and RBP target overlap” edges. Because these edge scores arise from different underlying statistics, they are not directly comparable across edge types. To enable interpretable visualization, both node and edge scores were categorized into four bins, where the mapping from raw RNA abundance and interaction significance values to binned node and edge scores is detailed in **Figure S3** and **Figure S4D**, respectively. In snoFlake, node scores are reflected by node size and edge scores by edge thickness.

snoFlake was generated using a Snakemake workflow^21^ available at https://github.com/scottgroup/snoFlake and visualized using Cytoscape (v3.10.1)^33^. All snoFlake node and edge annotations are provided in **Table S1** and an interactive Cytoscape session file is available as **Data S1**.

##### snoRNA and RBP target overlap analysis

To compute “snoRNA and RBP target overlap” edges, processed snoRNA–RNA and RBP–RNA interactions were used as input (**Figure S1**), restricting both to binding interactions on protein-coding RNA targets (Ensembl biotype: protein_coding)^62^ as they are the most consistently annotated across interaction datasets and are well suited for isoform-aware analyses. For each possible snoRNA–RBP pair in snoFlake, enrichment of overlapping binding sites on shared protein-coding RNA targets was assessed using bedtools fisher -s (v2.31.1)^76^ with the Ensembl GRCh38 primary assembly (release 110) as the reference genome, which tests whether the observed overlap exceeds that expected by chance given the genome-wide coverage of snoRNA and RBP binding sites (one-tailed Fisher’s exact test; see Quantification and Statistical Analysis for more details). Binding sites were considered overlapping if they shared ≥ 1 nt on the same strand.

##### Network motif analysis

A network motif was defined as a subgraph containing one snoRNA connected to two or more RBPs through double-edge interactions (snoRNA and RBP connected by both “RBP binds snoRNA” and “snoRNA and RBP target overlap” edges), with these RBPs additionally required to form a connected subgraph via “RBP binds RBP” edges (**Figure 2D**). Disconnected RBP groups linked to the same snoRNA were treated as separate motifs.

Motif enrichment was evaluated by comparing motif counts in snoFlake with those in 1,000 degree-preserving randomized networks generated using the double-edge swap algorithm. Motif counts included all occurrences of a given topology, such that overlapping motifs of lower complexity were counted independently. For each randomization, edges were iteratively rewired (10 swap attempts per edge) while preserving node degrees, node types, and interaction types. For each motif type, the *Z*-score was calculated as Z= (c_observed_ - µ_random_)/a_random_, where c_observed_ is the motif count in snoFlake and µ_random_ and a_random_ are the mean and standard deviation of motif counts across the 1,000 randomized networks. Empirical *p*-values were calculated and adjusted for multiple testing as described in Quantification and Statistical Analysis. Network randomization and motif analysis were performed using NetworkX (v3.4.2)^86^. Individual instances of each significantly enriched motif type are shown in **Figure S8**, displayed at the highest-order topology and colored by curated RBP function. The highest-scoring instance, determined by the sum of constituent node and edge scores, was selected for in-depth characterization as a candidate snoRNA–RBP regulatory complex.

#### Characterization of the SNORD22–PRPF8–EFTUD2 module

##### Structural modeling of the SNORD22–PRPF8–EFTUD2 complex

The secondary structure of SNORD22 was predicted using LinearPartition --mea^72^ and visualized using forna^87^ (**Figure 3B**). PRPF8 and EFTUD2 binding sites on SNORD22 were identified by intersecting processed ENCODE eCLIP datasets with SNORD22 genomic coordinates using bedtools intersect -s (v2.31.1)^76^. The primary target interaction region of SNORD22 was determined by aggregating all SNORD22 interactions that bind to protein-coding RNA targets and identifying nucleotide positions in the snoRNA sequence with high interaction density (**Figure S9**).

The three-dimensional structure of the SNORD22–PRPF8–EFTUD2 complex was modeled using the AlphaFold3 webserver^44^ with default parameters and visualized in PyMOL (v3.1.0)^73^ (**Figure 3C**). The SNORD22–PRPF8 interface was defined as amino acid–nucleotide contacts within 5 Å. No direct contacts were detected between SNORD22 and EFTUD2 within this distance threshold. The predicted complex was subsequently superimposed onto the cryo-EM structure of the human U5 snRNP (PDB: 8Q91)^45^ in PyMOL to evaluate the compatibility of the predicted SNORD22–PRPF8 interaction with the native U5 snRNP architecture (**Figure S10**).

##### RNA immunoprecipitation (RIP)

RNA immunoprecipitation (RIP) assays were performed in the ovarian serous cystadenocarcinoma cell line SKOV3.ip1 (RRID: CVCL_0C84)^50^ to assess RNA association with PRPF8 and EFTUD2. Cells (2 × 10^7^) were harvested by trypsinization and lysed in 150 µL ice-cold RIP lysis buffer containing 10 mM HEPES (pH 7.0), 100 mM KCl, 5 mM MgCl₂, 0.5% NP-40, 1 mM DTT, 0.25 U/mL RNase inhibitor (Protein Purification Platform, Université de Sherbrooke), protease inhibitor cocktail (1×; Complete, EDTA-free; Roche Diagnostics) and 0.4 mM vanadyl ribonucleoside complexes (VRC). Lysates were frozen overnight at −80°C, clarified by centrifugation, and protein concentration was determined using a BCA assay.

For RIP, 100 µL of lysate (10 µg/µL) was equilibrated for 1 h at room temperature and then incubated overnight at 4°C with rotation with 100 µL Dynabeads™ M-280 Sheep Anti-Rabbit IgG (Thermo Fisher Scientific, Cat# 11204D) pre-coupled with 5 µg of anti-PRPF8 (Proteintech, Cat# 111171-1-AP; RRID: AB_2171179), anti-EFTUD2 (Proteintech, Cat# 10208-1-AP; RRID: AB_2095834) or normal rabbit IgG (Cell Signaling Technology, Cat# 2729; RRID: AB_1031062). The immunoprecipitation was carried out in a final volume of 1 mL RIP buffer containing 100 mM NaCl, 50 mM Tris-HCl (pH 7.4), 0.04% NP-40, 0.2 U/mL RNaseOUT (Thermo Fisher Scientific, Cat# 10777019), 0.4 mM VRC, 1 mM DTT, 20 mM EDTA, and protease inhibitor cocktail (1:7). Parallel IgG RIPs were included as negative controls.

Beads were washed six times with NT2 wash buffer (150 mM NaCl, 50 mM Tris-HCl pH 7.4, 1 mM MgCl₂, 0.05% NP-40). After washing, beads were divided; one fraction (50%) was used for protein validation by western blot and the remaining fraction was processed for RNA extraction. Western blot data are representative of the RIP procedure but do not necessarily correspond to the same biological samples used for RIP–qPCR analysis (**Figure S11**).

##### Western blot validation of RIP

RIP and IgG control samples were eluted from beads by boiling at 95°C for 10 min in Laemmli buffer (62.5 mM Tris-HCl pH 6.8, 10% glycerol, 5% β-mercaptoethanol, 2.3% SDS). For input controls, 10 µL of lysate was collected prior to overnight antibody incubation, diluted in 1× Laemmli buffer, and denatured at 95°C for 5 min. Proteins were resolved by SDS–PAGE using 8% gels for PRPF8 and 10% gels for EFTUD2 and transferred to HyBond™ ECL membranes (Cytiva) for 2 h at 100 V.

Membranes were blocked overnight at 4°C in TBST (0.15 M NaCl, 0.02 M Tris-HCl pH 7.6, 0.1% Tween-20) supplemented with 5% skim milk, then incubated for 90 min at room temperature with agitation with the following primary antibodies: anti-PRPF8 (1:1500; Abcam, Cat# ab79237; RRID: AB_1925360), anti-EFTUD2 (1:2000; Proteintech, Cat# 10208-1-AP; RRID: AB_2095834), and anti-GAPDH (1:5000; Novus Biologicals, Cat# NB300-221; RRID: AB_350535). After washing, membranes were incubated for 90 min with HRP-conjugated secondary antibodies (1:2000; anti-rabbit IgG, Cytiva, Cat# NA934, RRID: AB_772206; anti-mouse IgG, Cytiva, Cat# NA931, RRID: AB_772210). Proteins were detected using Clarity™ Western ECL substrate (Bio-Rad) and imaged with a LAS-4000 system (Cytiva).

##### RNA isolation from RIP and RT–qPCR

RNA associated with immunoprecipitated ribonucleoprotein complexes was extracted by incubating magnetic beads in Proteinase K buffer (1% SDS, 1.2 mg/mL proteinase K, completed with NT2 buffer to a final volume of 150 µL) at 55°C for 30 min with shaking. RNA was purified by phenol–chloroform extraction, precipitated if needed, and resuspended in 10 µL of nuclease-free water. RNA concentration and purity were assessed using a NanoDrop spectrophotometer (Thermo Fisher Scientific). For reverse transcription, 7 µL of RNA was used per reaction and 2 µL was reserved for a no-reverse transcriptase (−RT) control to assess potential genomic DNA contamination. Reverse transcription was performed using gene-specific primers and the SuperScript IV Reverse Transcriptase kit (Thermo Fisher Scientific, Cat# 18091050) following the manufacturer’s protocol. Before qPCR, all cDNA samples were normalized to a final concentration of 3.33 ng/µL. Quantitative PCR was performed using gene-specific primers for SNORD22 and the reference gene RPPH1 (**Table S2**), with Power SYBR Green Master Mix (Thermo Fisher Scientific, Cat# 4368708) on a QuantStudio 6 Flex instrument (Applied Biosystems). Relative enrichment in RIP samples was calculated as 2*^ΔΔCq^,* where *ΔΔCq* = (*Cq_SNORD22 input_ - Cq_SNORD22 IP_*) - (*Cq_RPPH1 input_ - Cq_RPPH1 IP_*) and normalized to the IgG control. Each antibody (PRPF8, EFTUD2, and IgG) was assayed in four independent biological replicates (**Figure 3D**).

##### Binding profiles and sequence motif analyses

SNORD22, PRPF8, and EFTUD2 binding interactions were obtained and processed as described in Data Curation. Significance thresholds applied to each dataset are described in Quantification and Statistical Analysis.

To assess positional enrichment of SNORD22 binding interactions relative to the protein-coding transcriptome, binding windows were assigned to exonic, intronic, or exon-intron junction categories using the custom annotation file Homo_sapiens.GRCh38.110_snoRNAs_tRNAs.sorted.gtf. The distribution of SNORD22 binding windows across these categories was compared to the nucleotide composition of the protein-coding transcriptome, defined as the fraction of nucleotides in each category across all annotated exons, introns, and exon-intron junction positions (**Figure S12**).

To dissect overlapping binding patterns of SNORD22, PRPF8, and EFTUD2, interactions were categorized into mutually exclusive sets using bedtools intersect -s (v2.31.1)^76^ to identify co-binding regions, followed by bedtools subtract -s to isolate unique binding sites. This resulted in seven mutually exclusive categories of unique and co-binding interactions (**Figure S13**), with SNORD22 co-binding defined as the union of all categories containing both SNORD22 and at least one of PRPF8 or EFTUD2.

Transcriptome-wide binding profiles were generated from an intron-centric perspective comprising four regions: upstream exon, 150 nt window downstream of the 5′ splice site, 150 nt window upstream of the 3′ splice site, and the downstream exon (**Figures 3E and S15**). Binding scores represent normalized interaction density, calculated as (interaction count / number of features covering that position) × 10^7^, where features denote exons or the cumulative count of introns extending to each intronic position.

Enriched 6-mer sequence motifs^88^ were identified using MEME (v5.5.0)^48^ with the following parameters: -rna -mod zoops -minw 6 -maxw 6 -evt 0.05 -time 14400 - nmotifs 10 -objfun classic -markov_order 0. Motifs with E-values ≤ 1×10^−□^ were considered significant. Motif occurrences within binding profile regions were identified using FIMO (v5.5.0)^89^ with *p* ≤ 0.001.

##### Gene ontology enrichment analysis

Gene ontology (GO) enrichment analysis was performed using g:Profiler (*p*_adj_ < 0.01 and intersection size ≥ 10; version e113_eg59_p19_f6a03c19)^46^ on protein-coding RNA targets co-bound by SNORD22 and at least one of PRPF8 or EFTUD2. All protein-coding genes expressed in at least one of the five cancer cell lines (HCT116, MCF7, PC3, TOV112D, SKOV3.ip1)^23,24^ were used as background and GO terms for biological process (GO:BP), molecular function (GO:MF), and cellular component (GO:CC) were analyzed (**Figure S14**). Related terms were grouped by semantic similarity using GO-Figure! (similarity cutoff = 0.5)^47^.

#### SNORD22 knockdown analysis

##### RNA extraction and RT–qPCR

For knockdown experiments, SKOV3.ip1 cells were seeded at 300,000 cells per well in six-well plates. Four hours after seeding, cells were transfected with Lipofectamine 2000 (Invitrogen, Cat# 11668019) following the manufacturer’s protocols. Antisense oligonucleotides (ASOs) targeting SNORD22 (5-10-5 gapmers with 2′-O-methoxyethyl (2′MOE) bases; Integrated DNA Technologies) were used at a final concentration of 30 nM, with a non-targeting control ASO as negative control. Two ASOs were used to control for off-target effects. After 48 h, cells were harvested, and total RNA was extracted using the RNeasy Mini Kit (Qiagen, Germany) following the manufacturer’s guidelines, with the modification of adding 1.5 volumes of 100% ethanol during the precipitation step to preserve small RNAs. RNA quantity was measured using a NanoDrop spectrophotometer, and RNA integrity was assessed with the 2100 Bioanalyzer (Agilent Technologies, USA).

For RT–qPCR, 500 ng of RNA was used to validate knockdown. The reaction mixture and cycling conditions for reverse transcription and qPCR were performed as previously described^90^. Normalized relative expression was calculated using the ΔΔCq method, where Cq represents the quantification cycle. ASO sequences are provided in **Figure S16A** and primer sequences are listed in **Table S2**.

##### Preparation and sequencing of poly(A)-selected RNA-seq libraries

Each of the RNA-seq libraries was generated from 100 ng of total RNA free of genomic DNA using the NEBNext® Ultra™ II Directional RNA Library Prep Kit for Illumina (New England Biolabs, Cat# E7760S) and following the protocol for use with the NEBNext Poly(A) mRNA Magnetic Isolation Module (New England Biolabs, Cat# E7490). The resulting libraries were subjected to a total of 12 amplification cycles and then purified using 0.9× Ampure XP beads. Quality and size were assessed with an Agilent 2100 Bioanalyzer. The libraries were then quantified using a Qubit fluorometer, pooled at an equimolar concentration of 1 nM, then sequenced with 2 × 75 bp on an Aviti instrument from Element Biosciences using one lane of a Cloudbreak Freestyle High Output 150 cycles kit (Cat# 820-00022).

##### Differential expression and splicing analysis

Raw poly(A)-selected RNA-seq reads were processed using the Snakemake-based RNA-seq analysis workflow (https://github.com/kristinassong/RNAseq), which follows the same steps as the TGIRT-seq pipeline described above until STAR alignment (v2.7.11b). For poly(A)-selected RNA-seq analysis, only primary alignments from STAR were retained for further analyses using samtools view -b -F 256 (v1.22.1)^91^. Transcript abundances were quantified using kallisto quant --bias --bootstrap-samples=50 (v0.48.0)^92^ and coverage tracks were generated using bedtools genomecov -bg -split (v2.31.1)^76^.

Differential gene expression between SNORD22 knockdown (KD) and negative control (NC5) conditions was assessed using DESeq2 (v1.42.0)^51^ from four SNORD22 KD and two NC5 replicates. Genes were considered differentially expressed if they met the following criteria: protein-coding RNA abundance ≥ 1 TPM in at least one condition, |log_2_FC| > 1 and *p*_adj_ < 0.05. Differential alternative splicing analysis was performed using rMATS (v4.2.0)^52^. Significant alternative splicing events were required to have protein-coding RNA abundance ≥ 1 TPM in at least one condition, junction read count ≥ 10, |ΔPSI| ≥ 0.05 and FDR ≤ 0.01. Sashimi plots were generated using ggsashimi^93^ with the custom annotation defined in Data Curation.

##### Splice site strength analysis

The strength of the 5’ and 3’ splice sites within the flanking introns of cassette exons with decreased inclusion upon SNORD22 KD was evaluated using MaxEntScan^53^ (**Figure 4D**). Splice site sequences were extracted from the custom annotation defined in Data Curation using 3 nt in exon + 6 nt in intron for 5’ splice sites and 20 nt in intron + 3 nt in exon for 3’ splice sites. Scores were classified as weak (score < 8), moderate (8 ≤ score ≤ 9), or strong (score > 9) based on established thresholds^94^. A cassette exon was defined as having suboptimal splice site strength if either its upstream 3’ splice site or downstream 5’ splice site was weak.

##### Droplet digital PCR of SRRT and USP36

Droplet digital PCR (ddPCR) analysis was performed using 10 ng of cDNA per reaction, following the manufacturer’s protocol (Bio-Rad) and carried out by the Université de Sherbrooke RNomics Platform. The reaction conditions were as described in an earlier study^95^, with the exception of 50 PCR amplification cycles. Primer sequences are listed in **Table S2**.

##### Predicted functional consequences of exon exclusion

To investigate the predicted functional consequences of exon exclusion events following SNORD22 depletion, we used SpliceDecoder^74^. Events were classified according to the location of SNORD22 binding relative to the cassette exon: upstream intron splice sites, downstream intron splice sites, or intronic regions (mutually exclusive categories). Simulated exon exclusion events that did not correspond to annotated splice junctions were excluded. Predicted consequences were categorized as premature termination codon (PTC) removal, nonsense-mediated decay (NMD), gain of domain (GoD), loss of domain (LoD), altered UTR (UTR alt), altered CDS (CDS alt), or frame loss.

#### Quantification and statistical analysis

Statistical analyses were performed in Python (v3.11) and R (v4.3) unless otherwise noted.

##### Expression quantification and filtering

RNA abundance for snoRNAs and protein-coding transcripts was quantified from TGIRT-seq datasets as described above. SnoRNAs and RBPs with RNA abundance ≥ 1 TPM in at least one of the five cancer cell lines (HCT116, MCF7, PC3, TOV112D, SKOV3.ip1)^23,24^ were considered expressed and retained for downstream analyses.

##### Interaction filtering thresholds

For snoFlake network construction, snoGloBe interactions^32^ with scores ≥ 0.98 and HTRRI interactions^10,28–31^ supported by ≥ 5 chimeric reads were retained. ENCODE eCLIP interactions were filtered at *p* ≤ 10^−3^ and fold enrichment ≥ 8, following ENCODE guidelines^22^. STRING (v12.0)^27^ RBP–RBP interactions were filtered for physical links with a combined score ≥ 900, corresponding to high-confidence interactions.

For binding profile and SNORD22-specific analyses, less stringent thresholds were applied to increase coverage of SNORD22, PRPF8, and EFTUD2 binding sites: snoGloBe score ≥ 0.95, HTRRI ≥ 3 chimeric reads, and ENCODE eCLIP *p* ≤ 10^−2^, fold enrichment ≥ 4.

##### snoRNA and RBP target overlap significance

Overlap *p*-values from bedtools fisher (v2.31.1)^76^ were corrected for multiple testing using the Benjamini–Hochberg procedure to control the FDR < 0.001. The largest *p*-value satisfying the Benjamini–Hochberg criterion was identified and a significance threshold was set at the nearest more conservative order of magnitude. Consequently, all snoRNA–RBP pairs with *p* ≤ 10^−4^ were classified as having an enrichment of overlapping binding sites on shared protein-coding RNA targets and were represented by the “snoRNA and RBP target overlap” edge in snoFlake (**Figure 1A**).

##### Network motif enrichment

Empirical *p*-values for network motif enrichment were calculated as the fraction of randomized networks with motif counts greater than or equal to the observed count. *P*-values were adjusted for multiple testing using the Benjamini–Hochberg procedure. Motifs with *p*_adj_ < 0.05 were considered statistically significant.

##### RT–qPCR and ddPCR statistical analyses

Statistical comparisons for RT–qPCR and ddPCR experiments were performed using two-tailed *t*-tests. SNORD22 enrichment over IgG was compared between PRPF8 and EFTUD2 immunoprecipitates using Welch’s *t*-test on log ₂ - transformed enrichment values (*n* = 4 biological replicates; **Figure 3D**). SNORD22 KD efficiency was assessed by comparing each ASO KD sample against the negative control using a Student’s *t*-test (*n* = 2 biological replicates, 3 technical replicates each; **Figure S16C**). Exon inclusion levels from ddPCR splicing assays were similarly compared between each SNORD22 ASO KD sample and the negative control using a Student’s *t*-test (*n* = 3 biological replicates; **Figures S20B and S21B**).

##### Binding enrichment analysis

Enrichment of SNORD22 binding interactions at exonic, intronic, and exon-intron junction regions relative to the protein-coding transcriptome background was assessed using a chi-squared test of independence. Enrichment of SNORD22 co-binding with PRPF8 and/or EFTUD2 was assessed among differentially expressed genes vs. non-differentially expressed genes as well as cassette exons with increased or decreased inclusion vs. non-differentially spliced exons, using Fisher’s exact test.

##### Statistical analyses of exon exclusion events

Splice site strength score distributions and GC content were compared across groups using pairwise chi-squared tests. Differences in exon and intron lengths between significant and non-significant exon exclusion events were assessed using the Mann–Whitney *U* test. All were corrected for multiple testing using the Benjamini–Hochberg procedure. The frequency of PTC removal across SNORD22 binding location groups was compared using Fisher’s exact test.

