## Supplementary Figures for "snoFlake: A network model for snoRNA–RBP complexes reveals SNORD22 as a U5 snRNP-associated splicing regulator"

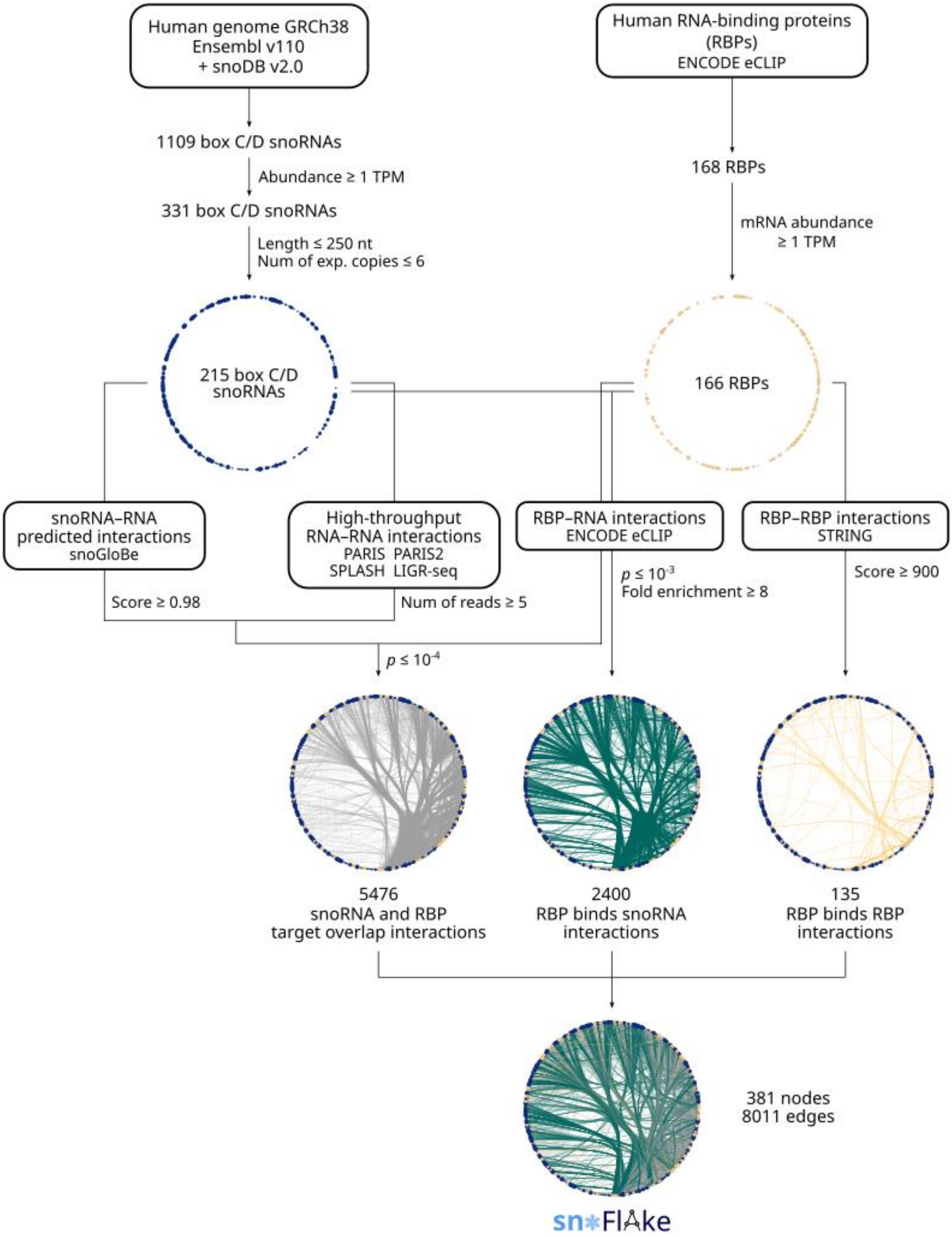

Human genome GRCh38
Ensembl v110
+ snoDB v2.0
Human RNA-binding proteins
(RBPs)
ENCODE eCLIP
1109 box C/D snoRNAs
Abundance ≥ 1 TPM
331 box C/D snoRNAs
Length ≤ 250 nt
Num of exp. copies ≤ 6
168 RBPs
mRNA abundance
≥ 1 TPM
215 box C/D
snoRNAs
166 RBPs
snoRNA–RNA
predicted interactions
snoGloBe
High-throughput
RNA–RNA interactions
PARIS PARIS2
SPLASH LIGR-seq
RBP–RNA interactions
ENCODE eCLIP
RBP–RBP interactions
STRING
Score ≥ 0.98
Num of reads ≥ 5
p ≤ 10-3
Fold enrichment ≥ 8
Score ≥ 900
p ≤ 10-4
5476
snoRNA and RBP
target overlap interactions
2400
RBP binds snoRNA
interactions
135
RBP binds RBP
interactions
381 nodes
8011 edges
sn*FlAke

**Figure S1. Overview of the snoFlake construction pipeline, related to Figure 1 and STAR Methods.** Human box C/D snoRNAs and RNA-binding proteins (RBPs) were first filtered based on RNA abundance, with additional filters applied to snoRNAs for sequence length and family copy number. Four complementary input datasets, including computationally predicted snoRNA–RNA interactions, high-throughput RNA–RNA interactions, RBP–RNA binding sites from ENCODE eCLIP datasets, and physical RBP–RBP interactions, were integrated to identify biologically meaningful snoRNA–RBP associations. Each intermediate network view highlights a distinct interaction layer, culminating in a comprehensive multidimensional interactome comprising 381 nodes and 8,011 edges, which together form the snoFlake network.

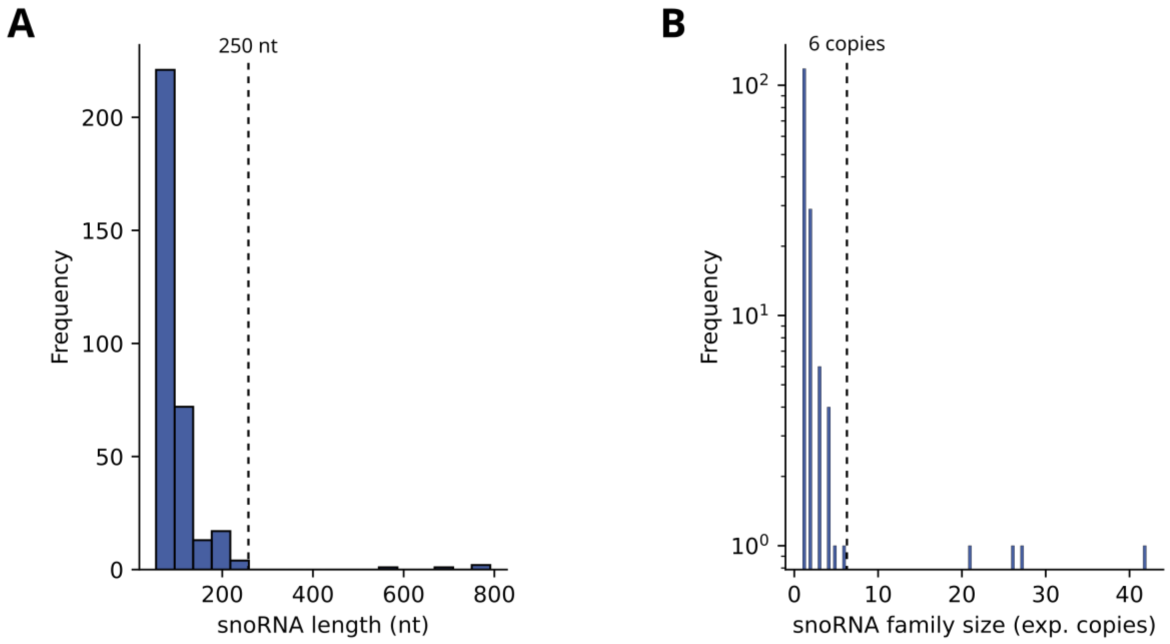


**Figure S2. Selection criteria for human box C/D snoRNAs included in snoFlake, related to Figure S1. (A)** Length distribution of all expressed human box C/D snoRNAs. Only snoRNAs with lengths ≤ 250 nt (left of the vertical dashed line) were retained to remain within the typical size range and ensure consistency across analyses. **(B)** Family size distribution of human box C/D snoRNAs based on Rfam, defined as the number of expressed copies per family. Only snoRNAs from families with ≤ 6 copies (left of the vertical dashed line) were included to exclude members of large, highly repeated families.

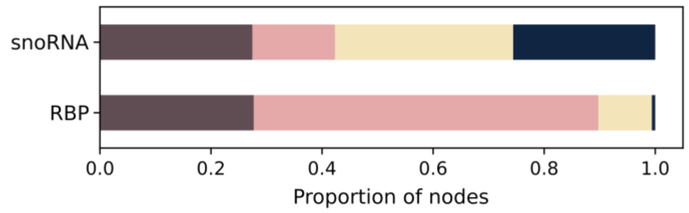


| | Score | RNA abundance |
|---|---|---|
| | 1 | 1–10 TPM |
| | 10 | 10–100 TPM |
| | 100 | 100–1000 TPM |
| | 1000 | ≥ 1000 TPM |

**Figure S3. Distribution of node scores in snoFlake, related to Figure 1B**. Proportional bar plot showing the distribution of node scores for box C/D snoRNAs and RBPs. Node scores were computed based on the maximum RNA abundance (in TPM) for each snoRNA and RBP across a compendium of five human cancer cell lines (HCT116, MCF7, PC3, TOV112D, and SKOV3.ip1) and assigned to one of four bins as shown in the table on the right.

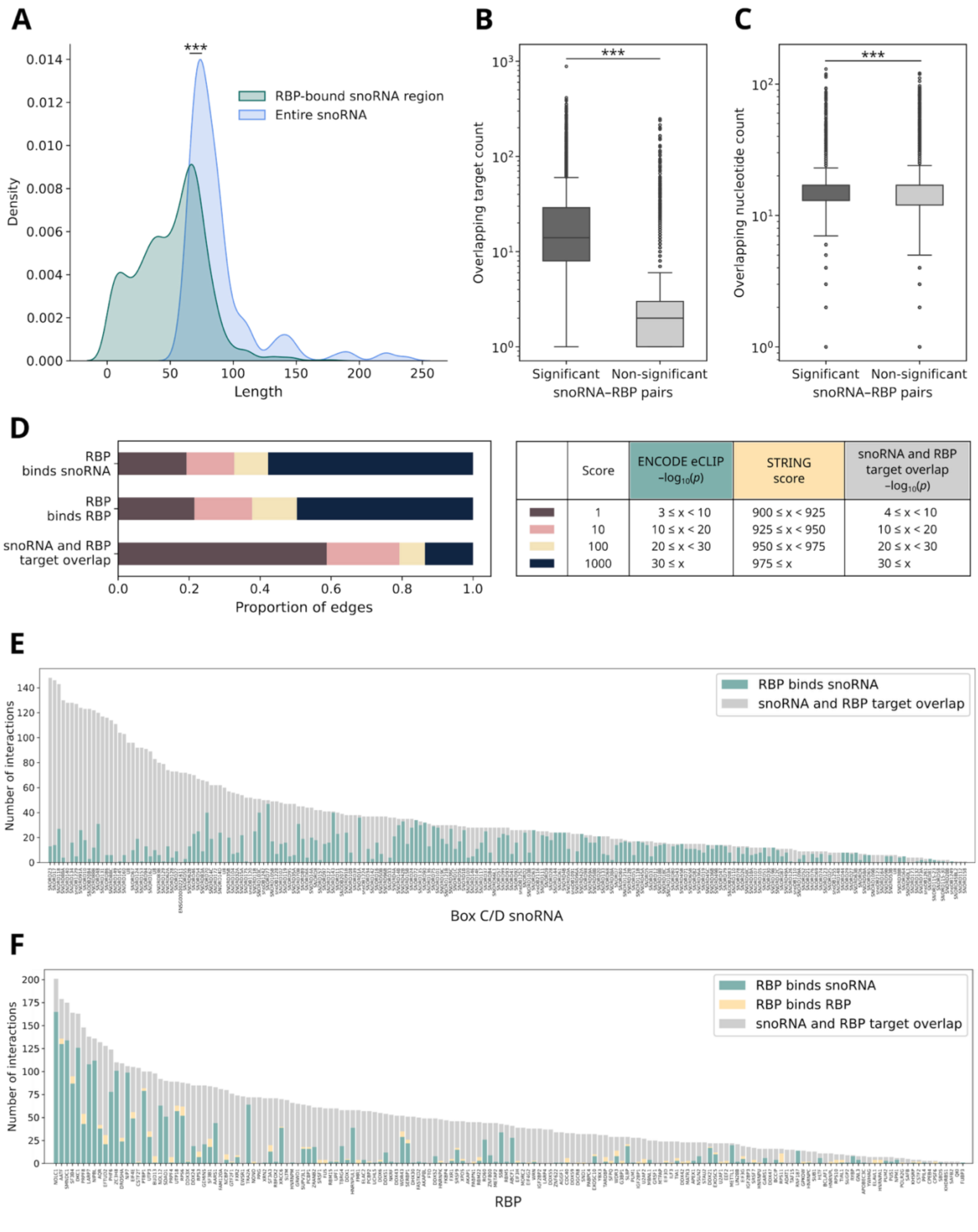
A
***
RBP-bound snoRNA region
Entire snoRNA
Density
Length
B
***
Overlapping target count
Significant
Non-significant
snoRNA–RBP pairs
C
***
Overlapping nucleotide count
Significant
Non-significant
snoRNA–RBP pairs
D
RBP binds snoRNA
RBP binds RBP
snoRNA and RBP target overlap
Proportion of edges
Score
ENCODE eCLIP −log10(p)
STRING score
snoRNA and RBP target overlap −log10(p)
1 | 3 ≤ x < 10 | 900 ≤ x < 925 | 4 ≤ x < 10
10 | 10 ≤ x < 20 | 925 ≤ x < 950 | 10 ≤ x < 20
100 | 20 ≤ x < 30 | 950 ≤ x < 975 | 20 ≤ x < 30
1000 | 30 ≤ x | 975 ≤ x | 30 ≤ x
E
RBP binds snoRNA
snoRNA and RBP target overlap
Number of interactions
Box C/D snoRNA
F
RBP binds snoRNA
RBP binds RBP
snoRNA and RBP target overlap
Number of interactions
RBP

**Figure S4. Comparative analyses of snoRNA and RBP interaction features in snoFlake, related to Figure 1A**. **(A)** Density plot comparing the total length distribution of box C/D snoRNAs with the length of snoRNA regions bound by an RBP based on ENCODE eCLIP data. Statistical significance was assessed using the Mann–Whitney *U* test (***$p < 0.001$). **(B)** Box plot showing the number of protein-coding RNA targets on which a snoRNA and an RBP co-bind at overlapping sites. Significant snoRNA–RBP pairs (Fisher's exact test, $p \leq 10^{-4}$; STAR Methods) share more overlapping targets than non-significant pairs. Statistical significance was assessed using the Mann–Whitney *U* test (***$p < 0.001$). **(C)** Same as (B), but comparing the length of individual co-bound sites on protein-coding RNA targets. **(D)** Proportional bar plot of edge scores for each interaction type in snoFlake. Edge scores reflect interaction significance, with score definitions provided in the table on the right. Table column headers are colored by the representative color of each interaction type: "RBP binds snoRNA" in green, "RBP binds RBP" in orange, and "snoRNA and RBP target overlap" in gray. **(E)** Stacked bar plot showing the number of interactions per snoRNA, categorized by interaction type. **(F)** Same as (E), but for RBPs.

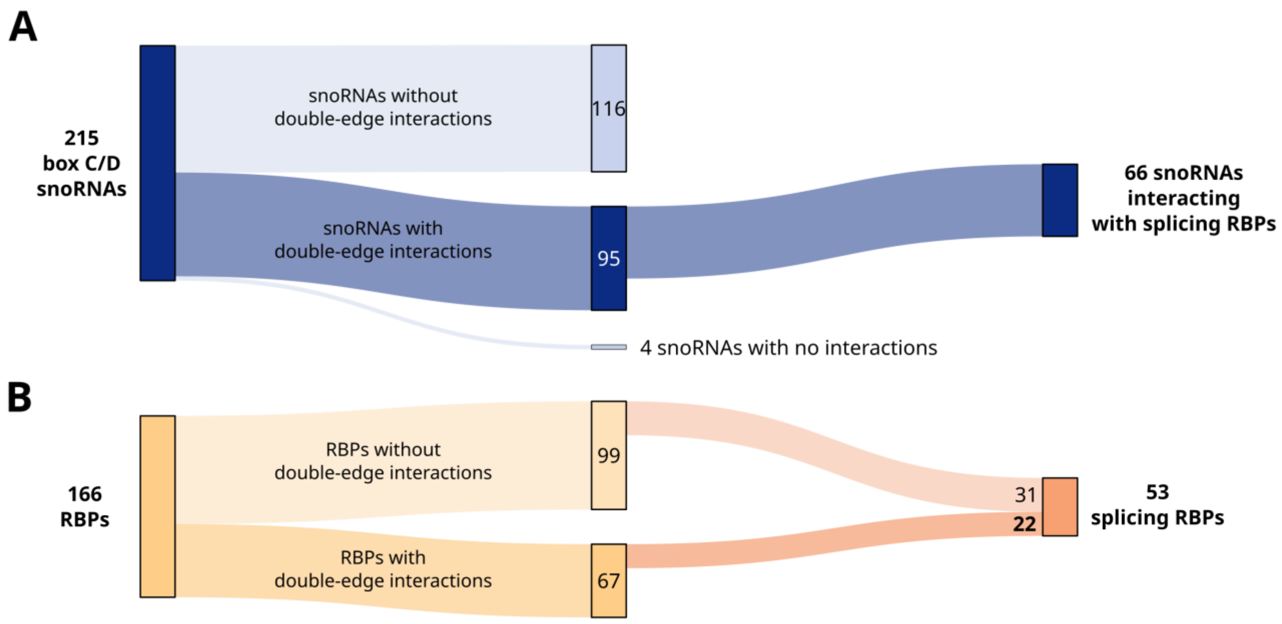


**Figure S5. Double-edge snoRNAs are frequently associated with a subset of splicing-related RBPs, related to Figure 2C.** Sankey plots showing **(A)** the number of snoRNAs and **(B)** the number of RBPs in snoFlake, split by whether they are involved in double-edge interactions. The subset of splicing-related RBPs involved in double-edge interactions and the snoRNAs that interact with them are highlighted in bold.

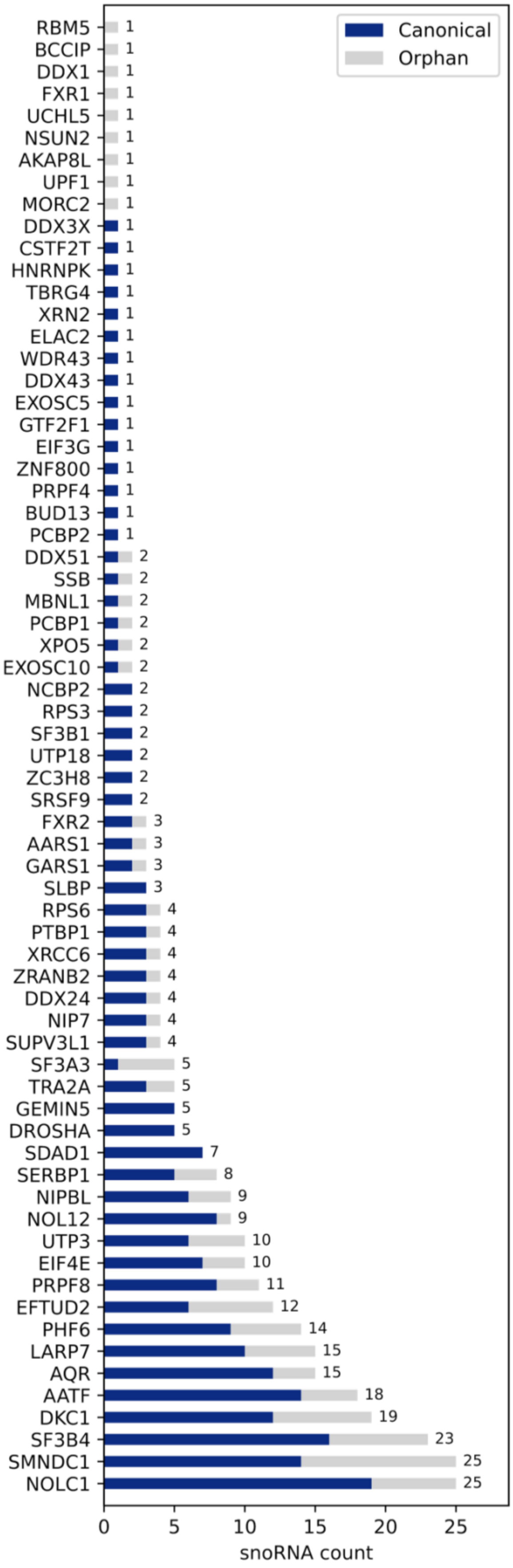
Canonical
Orphan
RBM5 1
BCCIP 1
DDX1 1
FXR1 1
UCHL5 1
NSUN2 1
AKAP8L 1
UPF1 1
MORC2 1
DDX3X 1
CSTF2T 1
HNRNPK 1
TBRG4 1
XRN2 1
ELAC2 1
WDR43 1
DDX43 1
EXOSC5 1
GTF2F1 1
EIF3G 1
ZNF800 1
PRPF4 1
BUD13 1
PCBP2 1
DDX51 2
SSB 2
MBNL1 2
PCBP1 2
XPO5 2
EXOSC10 2
NCBP2 2
RPS3 2
SF3B1 2
UTP18 2
ZC3H8 2
SRSF9 2
FXR2 3
AARS1 3
GARS1 3
SLBP 3
RPS6 4
PTBP1 4
XRCC6 4
ZRANB2 4
DDX24 4
NIP7 4
SUPV3L1 4
SF3A3 5
TRA2A 5
GEMIN5 5
DROSHA 5
SDAD1 7
SERBP1 8
NIPBL 9
NOL12 9
UTP3 10
EIF4E 10
PRPF8 11
EFTUD2 12
PHF6 14
LARP7 15
AQR 15
AATF 18
DKC1 19
SF3B4 23
SMNDC1 25
NOLC1 25
0
5
10
15
20
25
snoRNA count

**Figure S6. Double-edge snoRNA–RBP interactions span both canonical and orphan box C/D snoRNAs.** Bar plot showing the number of double-edge box C/D snoRNA interactors for each RBP shown on the y-axis. Bars are segmented and colored according to whether the interacting snoRNAs have known canonical modification targets (rRNA or snRNA; dark blue) or are orphan snoRNAs without known targets (gray). The total number of snoRNA interactors per RBP is indicated to the right of each bar.

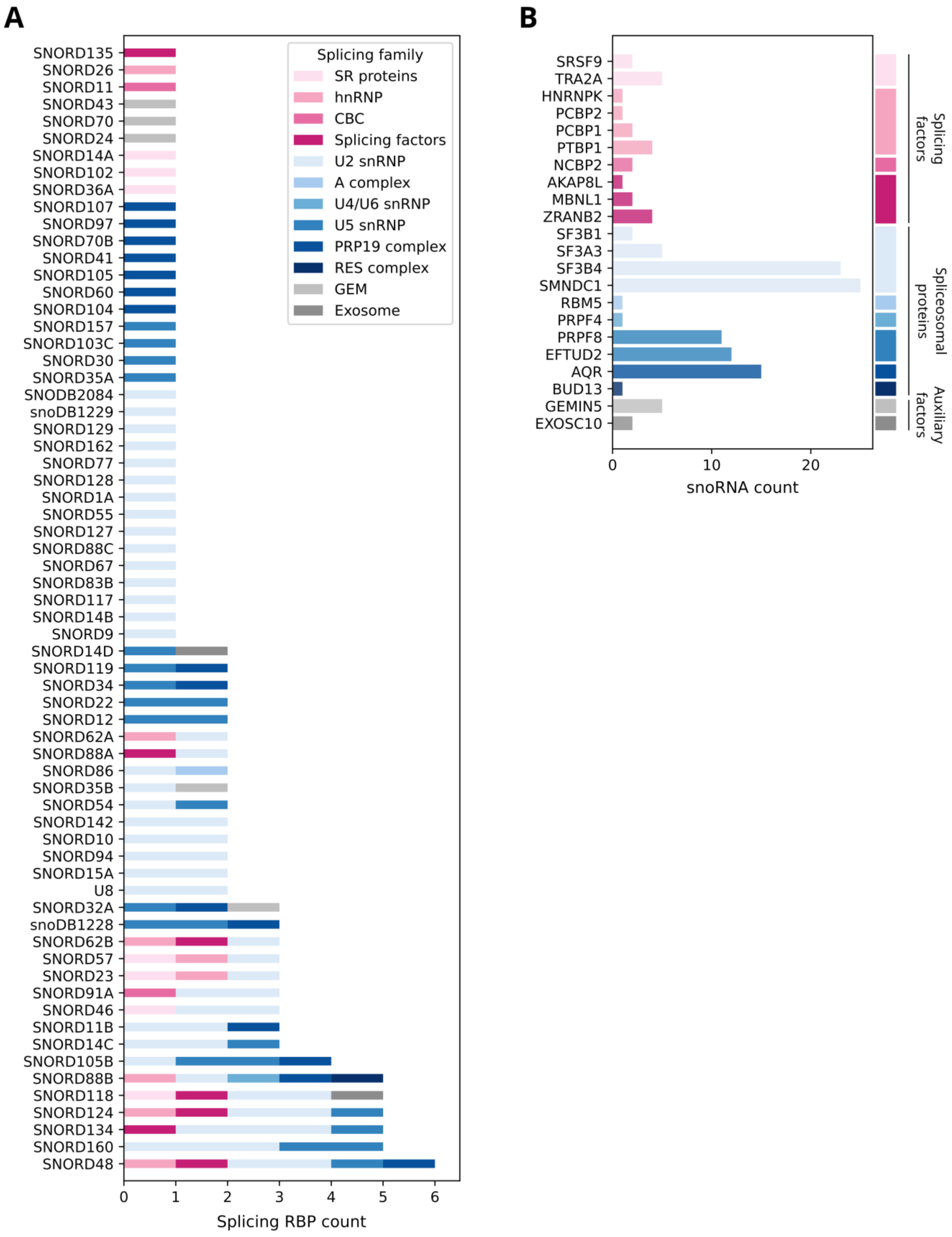
A
B
Splicing family
SR proteins
hnRNP
CBC
Splicing factors
U2 snRNP
A complex
U4/U6 snRNP
U5 snRNP
PRP19 complex
RES complex
GEM
Exosome
Splicing RBP count
snoRNA count
Splicing factors
Spliceosomal proteins
Auxiliary factors

**Figure S7. Splicing-related double-edges are dominated by recurring spliceosomal proteins, related to Figure S5. (A)** Bar plot showing the number of splicing-related RBPs involved in a double-edge interaction with each snoRNA on the y-axis. Bars are colored by the splicing family annotation of each RBP interactor: splicing factors in pink tones, spliceosomal proteins in blue tones, and auxiliary factors in gray tones. **(B)** Bar plot showing the number of double-edge snoRNA interactors per splicing-related RBP on the y-axis. Bars are colored according to the same scheme as in (A).

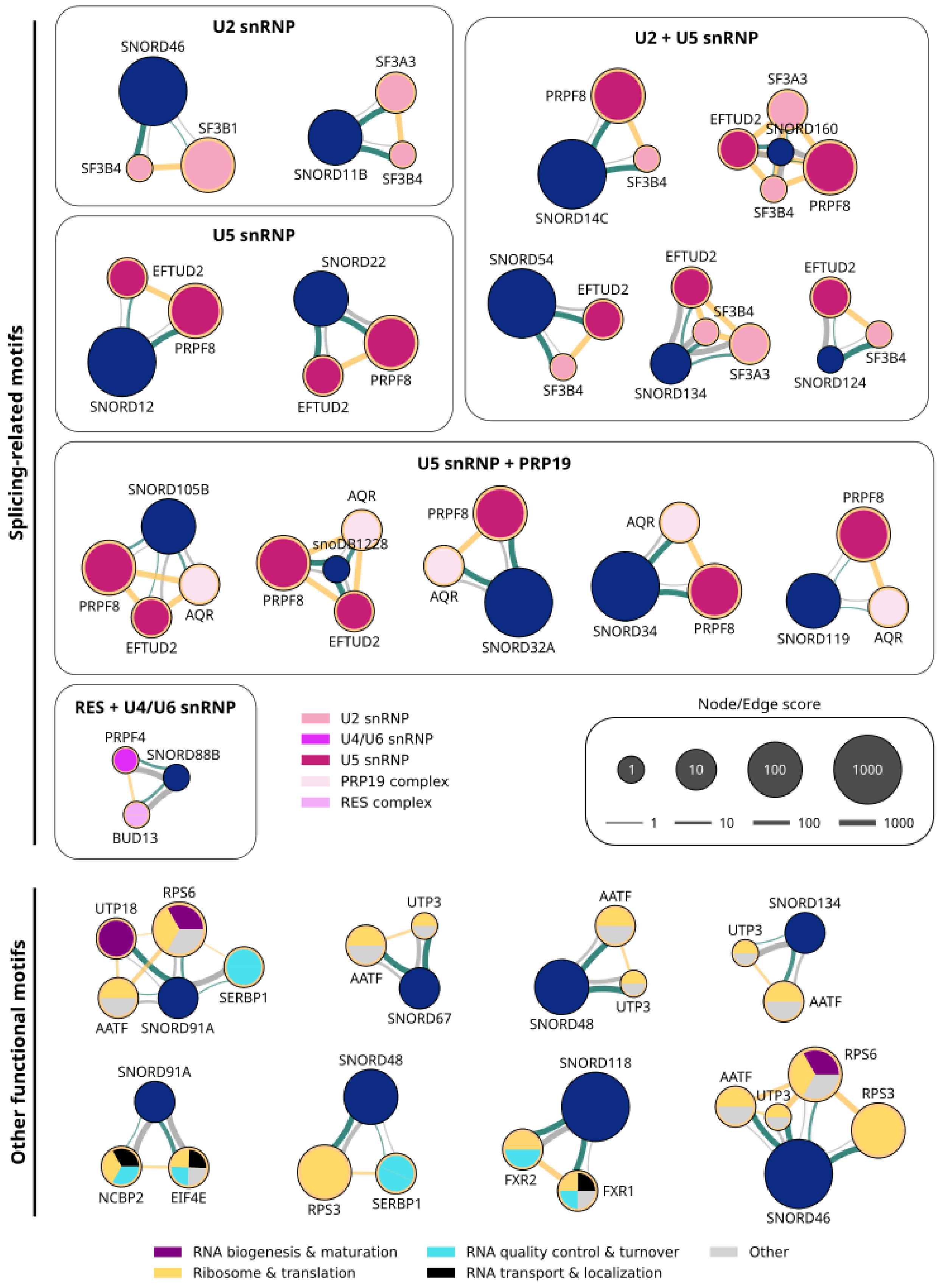
Splicing-related motifs
U2 snRNP
SNORD46
SF3B1
SF3B4
SF3A3
SNORD11B
SF3B4
U2 + U5 snRNP
PRPF8
SNORD14C
SF3B4
SF3A3
EFTUD2
SNORD160
SF3B4
PRPF8
SNORD54
EFTUD2
SF3B4
EFTUD2
SF3B4
SNORD134
SF3A3
EFTUD2
SF3B4
SNORD124
U5 snRNP
EFTUD2
PRPF8
SNORD12
SNORD22
PRPF8
EFTUD2
U5 snRNP + PRP19
SNORD105B
PRPF8
EFTUD2
AQR
AQR
snoDB1228
PRPF8
EFTUD2
PRPF8
AQR
SNORD32A
AQR
SNORD34
PRPF8
PRPF8
SNORD119
AQR
RES + U4/U6 snRNP
PRPF4
SNORD88B
BUD13
U2 snRNP
U4/U6 snRNP
U5 snRNP
PRP19 complex
RES complex
Node/Edge score
1
10
100
1000
1
10
100
1000
Other functional motifs
UTP18
RPS6
SERBP1
AATF
SNORD91A
UTP3
AATF
SNORD67
AATF
SNORD48
UTP3
SNORD134
UTP3
AATF
SNORD91A
NCBP2
EIF4E
SNORD48
RPS3
SERBP1
SNORD118
FXR2
FXR1
AATF
UTP3
RPS6
RPS3
SNORD46
RNA biogenesis & maturation
Ribosome & translation
RNA quality control & turnover
RNA transport & localization
Other

**Figure S8. Enriched snoRNA–RBP network motifs reveal modular spliceosomal assemblies, related to Figure 2D**. Individual instances of each significant motif type identified in Figure 2D are displayed at the highest-order topology observed for each snoRNA–RBP configuration. When a snoRNA forms motifs with more than one disconnected RBP cluster, each cluster is shown as a separate motif and the same snoRNA may therefore appear in multiple motifs. Node size and edge thickness are scaled by their respective scores, representing RNA abundance and interaction significance. Motifs are separated into splicing-related motifs and other functional motifs based on the functional annotation of the RBPs. For splicing-related motifs, RBPs are colored by splicing family annotation, and motifs sharing the same annotation composition are grouped together and labeled by their representative complex. For other functional motifs, RBPs are colored according to their annotated function, with multifunctional RBPs colored proportionally to reflect their functional diversity.

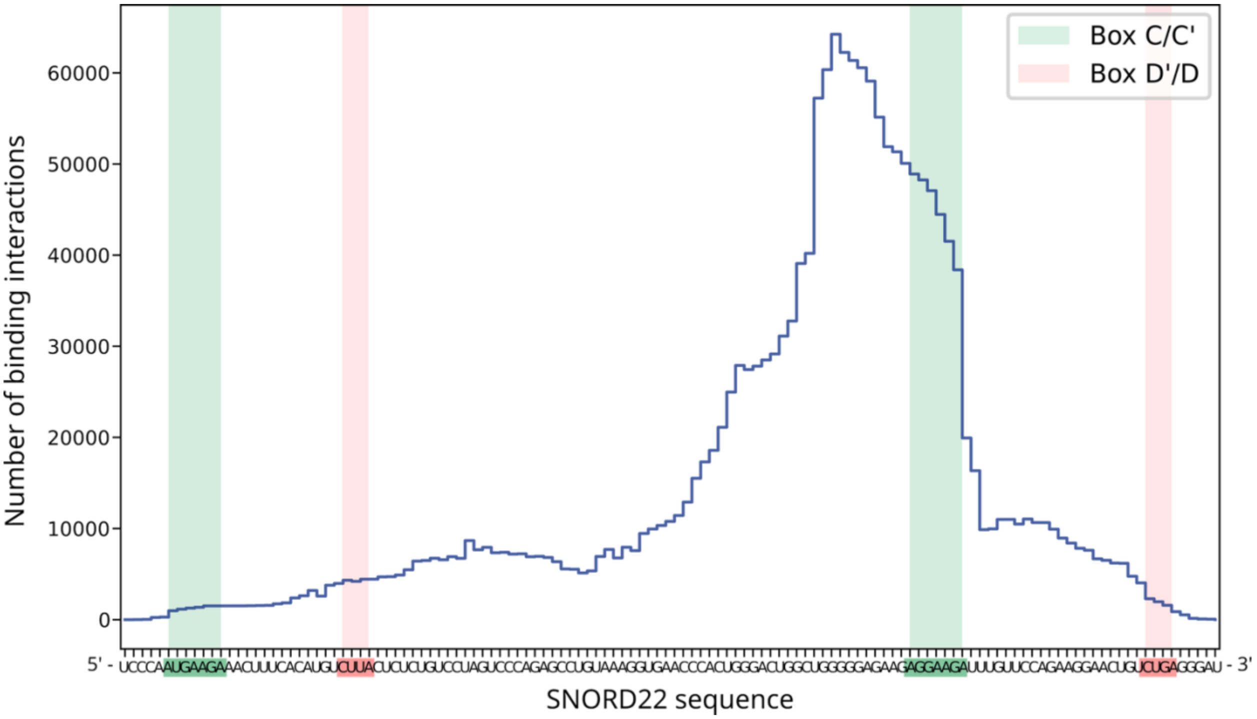


**Figure S9. SNORD22 uses a noncanonical region to bind to its protein-coding RNA targets, related to Figure 3B.** Line plot showing the number of predicted binding interactions per nucleotide along the SNORD22 sequence. The SNORD22 nucleotide positions are shown on the x-axis with the positions of the box C/C’ and D/D’ motifs highlighted in green and red, respectively.

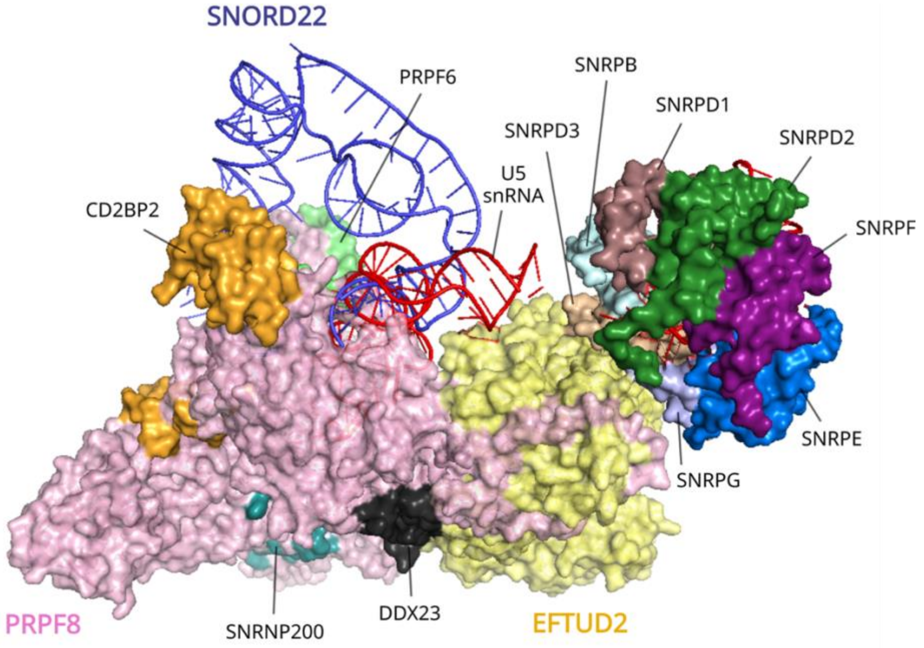


**Figure S10. Predicted SNORD22 interaction within the U5 snRNP structure, related to Figure 3C.** The AlphaFold3-predicted SNORD22–PRPF8–EFTUD2 complex is aligned to the cryo-EM structure of the human U5 snRNP (PDB: 8Q91) to assess the structural compatibility of the predicted interaction within the spliceosomal particle.

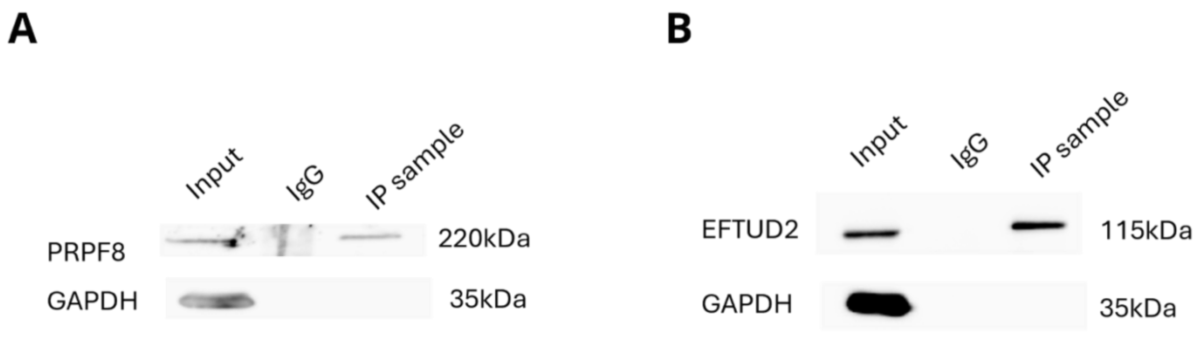


**Figure S11. Western blot validation of PRPF8 and EFTUD2 immunoprecipitation, related to Figure 3D. (A)** Representative western blot showing PRPF8 enrichment in the PRPF8 IP compared with the IgG control IP and input lysate. GAPDH was probed as a non-specific control. Molecular weights are indicated on the right. **(B)** Same as (A), but for EFTUD2.

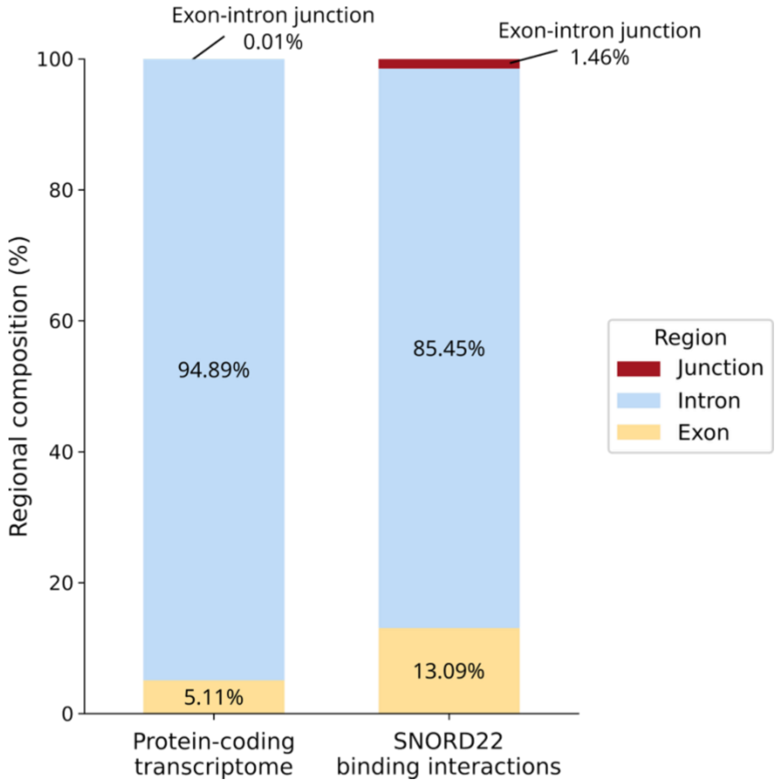


**Figure S12. SNORD22 binding interactions are enriched at exon-intron junctions and exons, related to Figure 3E.** Bar plot showing the proportion of nucleotides located in exon, intron and exon-intron junction regions in the protein-coding transcriptome compared to SNORD22 binding regions. For the protein-coding transcriptome, percentages are calculated as the fraction of nucleotides in each category relative to all nucleotides in exons, introns and annotated exon-intron junction positions. For SNORD22, percentages are calculated as the fraction of binding windows assigned to each category relative to all SNORD22 binding windows. Differences in regional composition between the protein-coding transcriptome and SNORD22 binding windows were assessed using a chi-squared test of independence ($\chi^2(2) \approx 5.0 \times 10^5$, $p < 10^{-16}$).

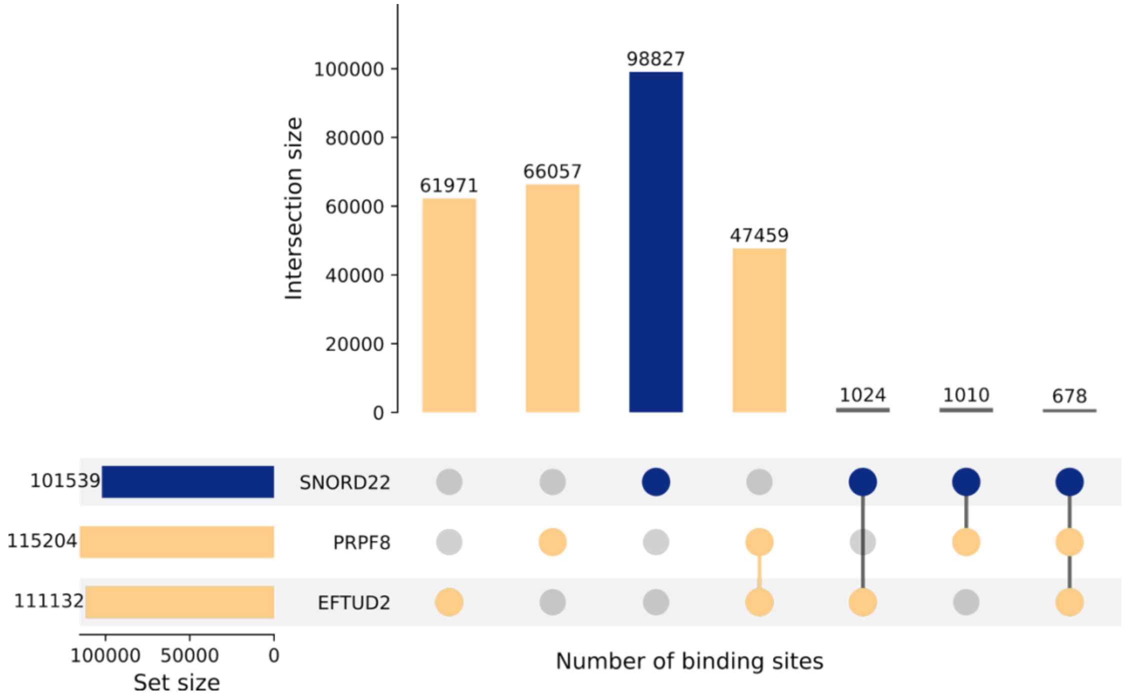


**Figure S13. Intersection of SNORD22, PRPF8 and EFTUD2 binding interactions, related to Figure 3E.** UpSet plot showing the overlap of SNORD22, PRPF8 and EFTUD2 binding interactions across protein-coding RNA targets in the transcriptome.

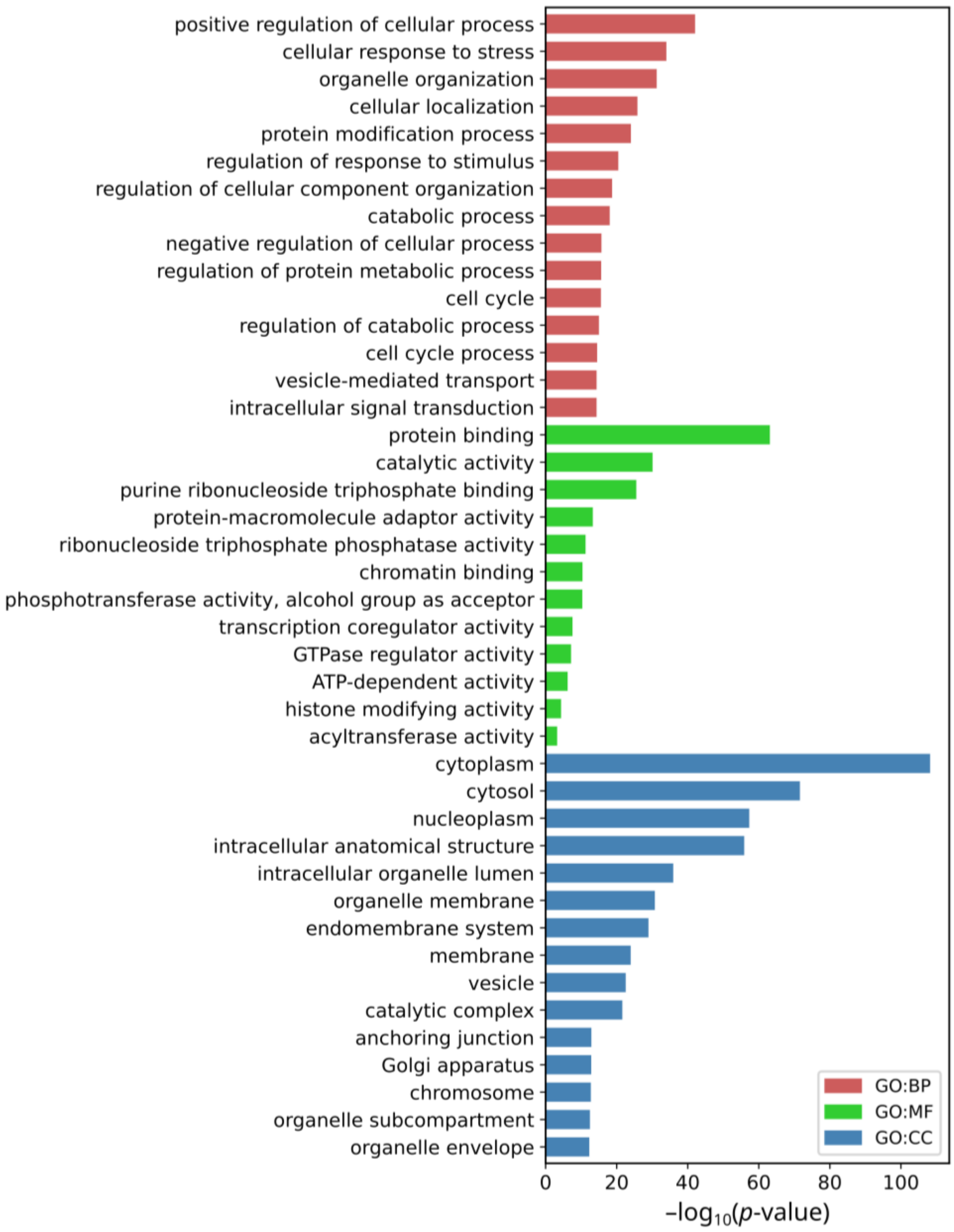


**Figure S14. Functional enrichment of protein-coding RNA targets co-bound by SNORD22 and PRPF8/EFTUD2, related to Figure S13.** Bar plot showing enriched Gene Ontology (GO) terms identified using g:Profiler ($p_{adj} < 0.01$ and intersection size ≥ 10) for biological process (GO:BP), molecular function (GO:MF) and cellular component (GO:CC). GO enrichment analysis was performed on the overlapping protein-coding RNA targets co-bound by SNORD22 and at least one of PRPF8 or EFTUD2 as defined in Figure S13. Similar GO terms were clustered based on semantic similarity using GO-Figure! (similarity cutoff = 0.5) and the top 15 driver terms for each ontology are shown on the plot.

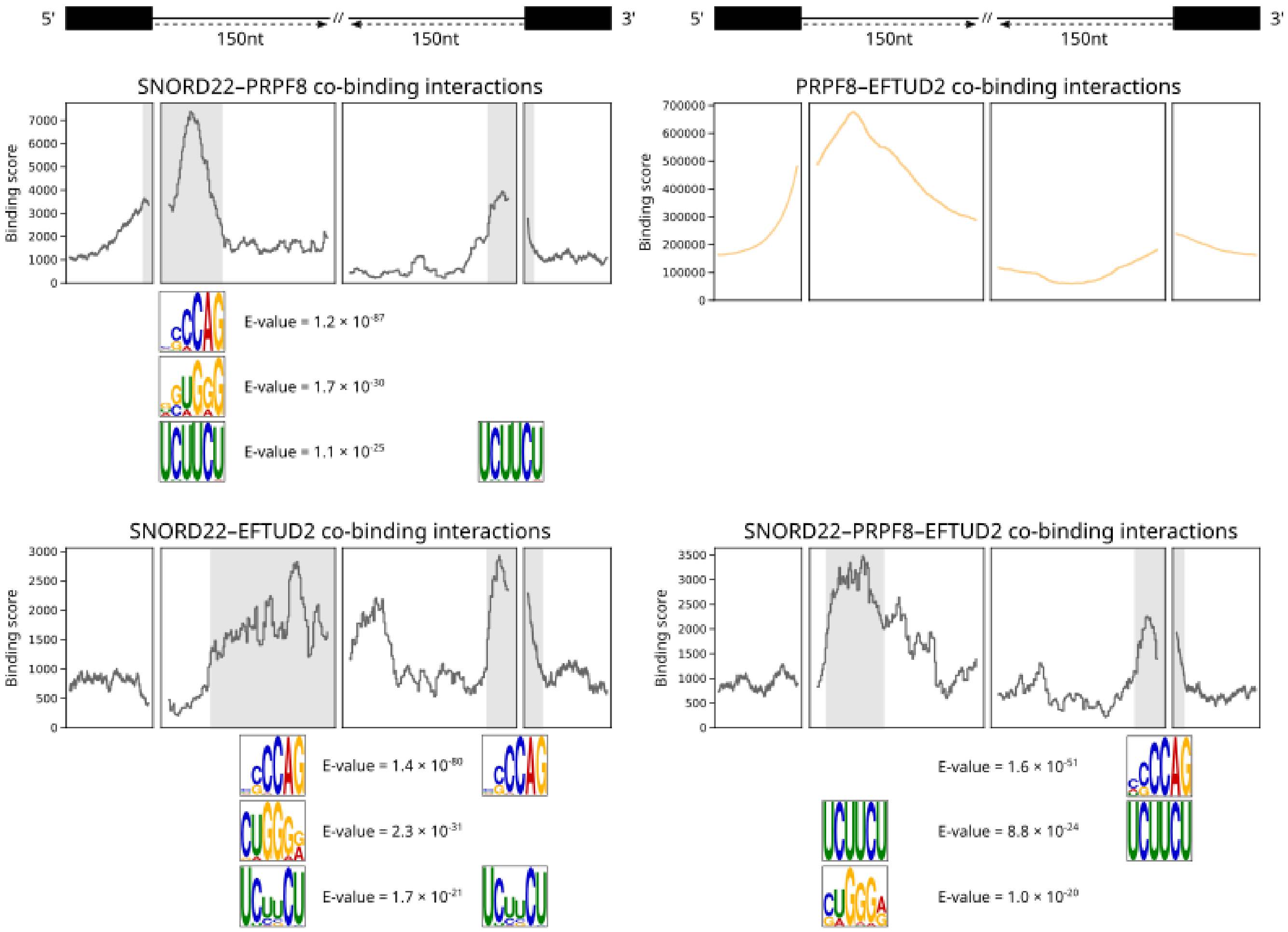


**Figure S15. Transcriptome-wide binding profiles of overlapping SNORD22, PRPF8 and EFTUD2 interactions, related to Figure 3E.** Binding profiles are shown for four mutually exclusive overlapping interaction categories: SNORD22–PRPF8, SNORD22–EFTUD2, PRPF8–EFTUD2 and SNORD22–PRPF8–EFTUD2. For each category, binding interactions on protein-coding RNA targets are collapsed and regions spanning splice sites (±150 nt around the two flanking exons) are shown. Peaks in the profiles indicate binding enrichment with the binding score shown on the y-axis. Significant binding motifs (MEME E-value ≤ $1\times10^{-5}$) identified using MEME are shown below each profile at the regions where they are enriched, together with their E-values.

**A**

| ASO | Sequence |
|---|---|
| SNORD22 ASO1 | 5' - mU*mA*mG*mG*mA*C*A*G*A*G*A*G*T*A*A*mG*mA*mC*mA*mU - 3' |
| SNORD22 ASO2 | 5' - mA*mU*mC*mC*mC*T*C*A*G*A*C*A*G*T*T*mC*mC*mU*mU*mC - 3' |
| NC5 | 5' - mG*mC*mG*mA*mC*T*A*T*A*C*G*C*G*C*A*mA*mU*mA*mU*mG - 3' |

**B** **C**

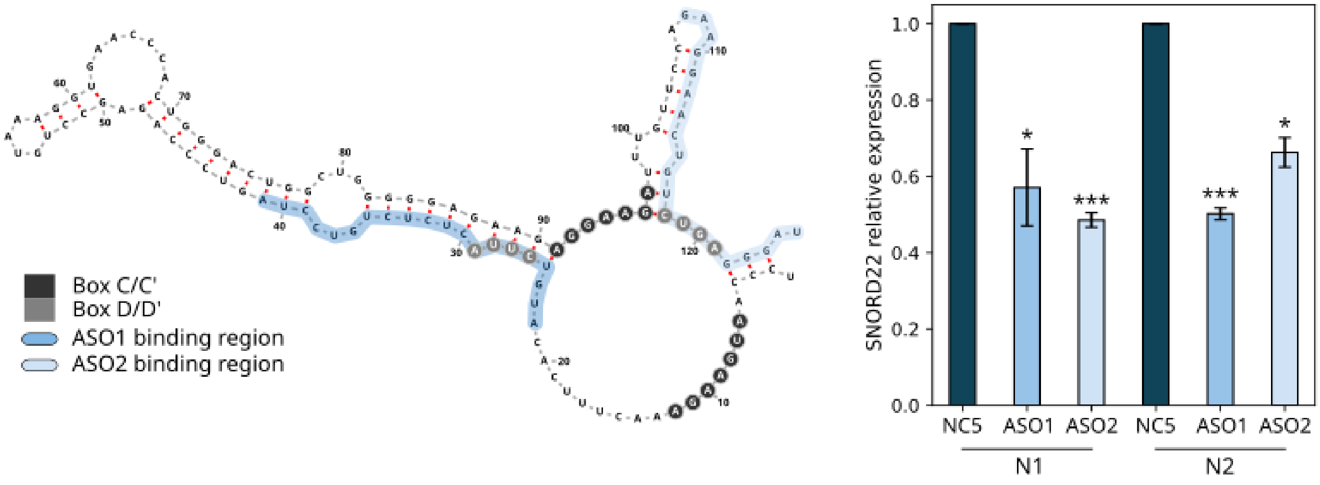


**Figure S16. SNORD22 knockdown using two different antisense oligonucleotides, related to Figure 4A. (A)** ASO sequences used for SNORD22 knockdown and a non-targeting control ASO (NC5) in SKOV3.ip1 cells. * Indicates phosphorothioate linkages and *m* indicates 2'-O-methoxyethyl-modified nucleotides. **(B)** Binding positions of the two ASOs mapped onto the predicted secondary structure of SNORD22. **(C)** Bar plot showing the relative expression level of SNORD22 measured by RT-qPCR after knockdown with each ASO (two biological replicates per condition). Error bars represent the standard deviation of three technical replicates. Statistical significance was assessed using a t-test (*$p < 0.05$; *** $p < 0.001$).

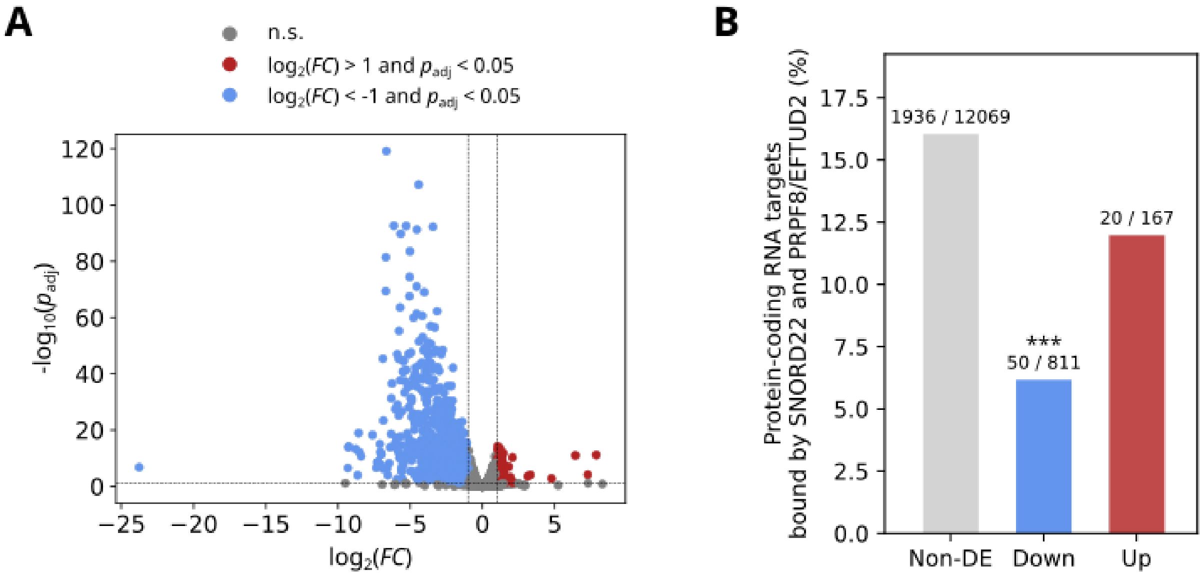


**Figure S17. Differentially expressed genes upon SNORD22 KD are not enriched for co-binding by SNORD22 and PRPF8/EFTUD2.** **(A)** Volcano plot of differentially expressed protein-coding RNA targets upon SNORD22 KD versus negative control NC5 (RNA abundance ≥ 1 TPM in at least one condition, $|\log_2 FC| > 1$ and $p_{adj} < 0.05$) identified using DESeq2. Downregulated ($n$ = 811) and upregulated ($n$ = 167) targets are shown in blue and red respectively. **(B)** Proportion of protein-coding RNA targets co-bound by SNORD22 and at least one of PRPF8 or EFTUD2 at the same binding site among non-differentially expressed (Non-DE), downregulated (Down) and upregulated (Up) genes defined in (A). Bars are labelled with the number of co-bound genes and the total number of genes in each category. Statistical significance was assessed using two-sided Fisher's exact tests comparing Down vs Non-DE and Up vs Non-DE (***$p < 0.001$).

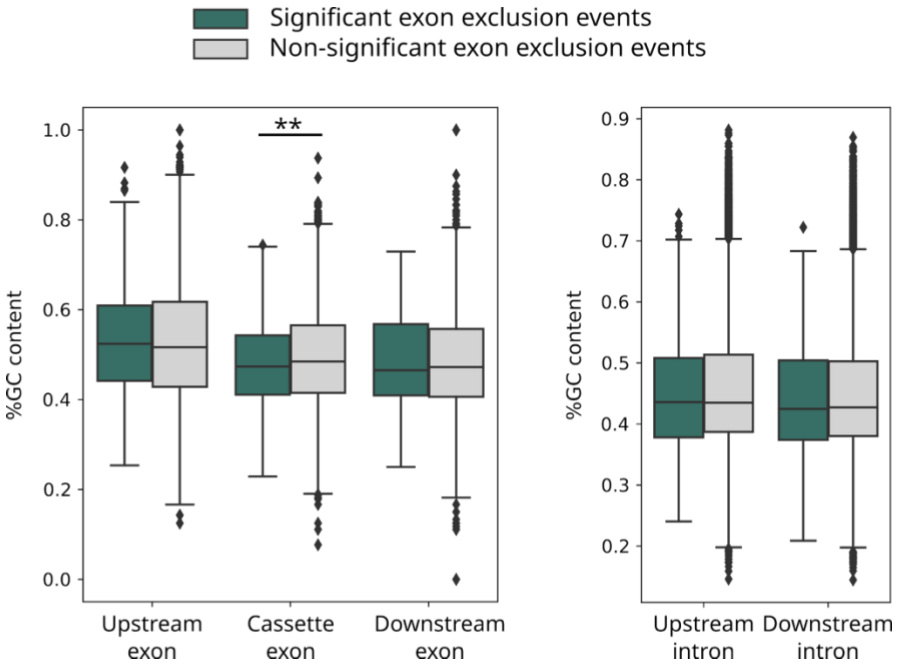


**Figure S18. Exons excluded upon SNORD22 KD have lower GC content, related to Figures 4E and 4F**. Box plots of exon and intron GC content for exon exclusion events upon SNORD22 KD. GC content is shown for the upstream, cassette and downstream exons as well as the upstream and downstream introns flanking the cassette exon. Distributions are shown separately for significant (|ΔPSI| ≥ 0.05 and FDR ≤ 0.01; dark green) and non-significant (|ΔPSI| < 0.05 or FDR > 0.01; gray) exon exclusion events. Pairwise chi-squared tests were used to compare GC content between significant and non-significant events for each exon and intron category (**$p_{adj} < 0.01$).

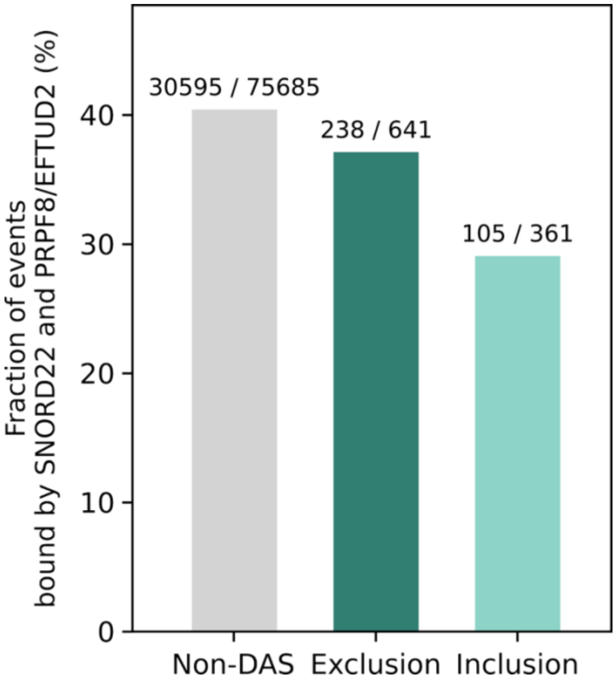


**Figure S19. Exon exclusion events are not globally enriched for co-binding by SNORD22 and PRPF8/EFTUD2, related to Figure 4G**. Bar plot showing the proportion of cassette exon events that are co-bound by SNORD22 and at least one of PRPF8 or EFTUD2 at the same binding site among all non-differentially spliced cassette exon events (Non-DAS), significant exon exclusion events (Exclusion) and significant exon inclusion events (Inclusion) upon SNORD22 KD as defined in Figure 4B. Bars are labelled with the number of co-bound events and the total number of events in each category. Statistical significance was assessed using two-sided Fisher's exact tests, comparing Exclusion vs Non-DAS ($p$ = 0.097) and Inclusion vs Non-DAS ($p = 9.56 \times 10^{-6}$).

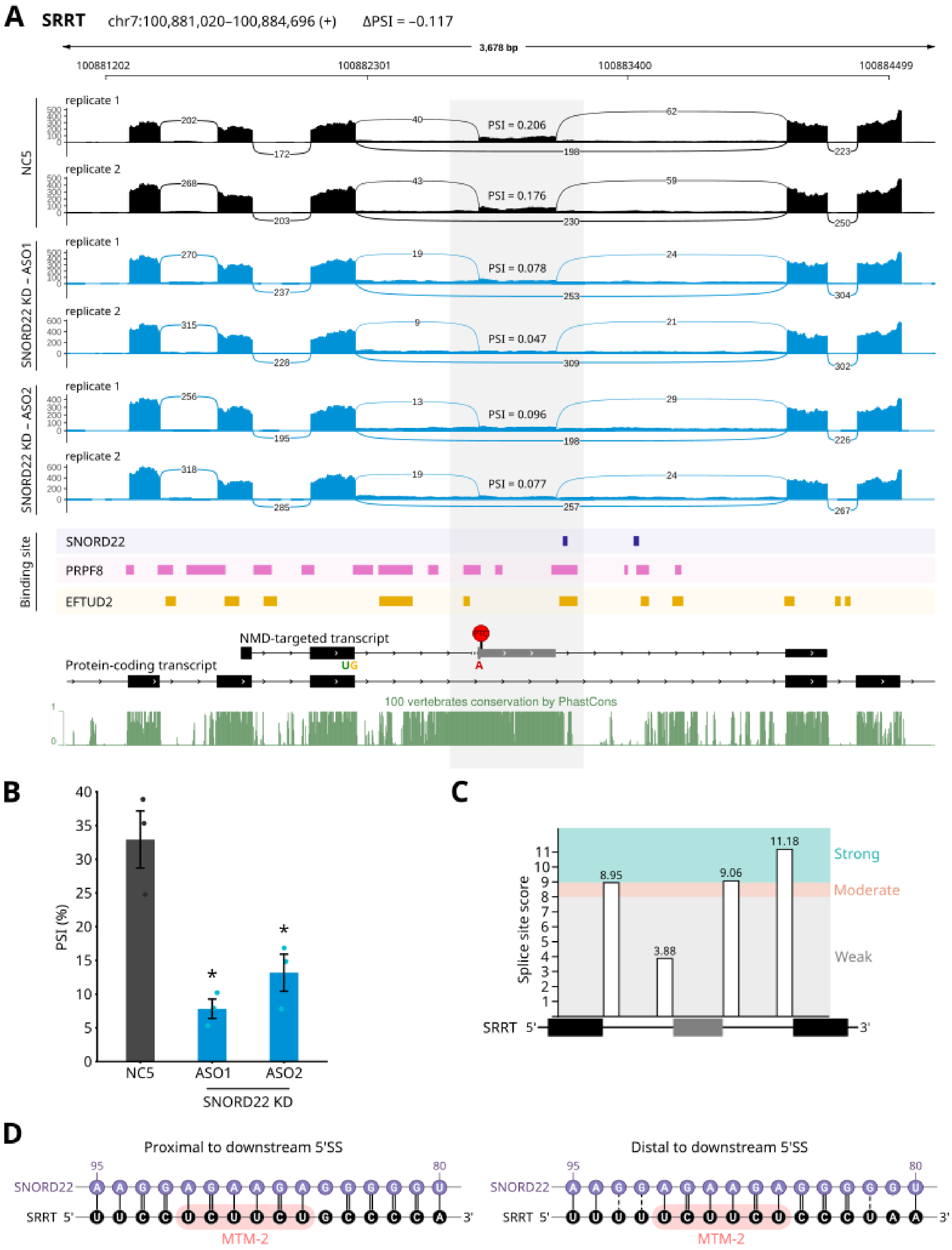
A
SRRT
chr7:100,881,020–100,884,696 (+)
ΔPSI = –0.117
3,678 bp
100881202
100882301
100883400
100884499
NC5
replicate 1
PSI = 0.206
replicate 2
PSI = 0.176
SNORD22 KD – ASO1
replicate 1
PSI = 0.078
replicate 2
PSI = 0.047
SNORD22 KD – ASO2
replicate 1
PSI = 0.096
replicate 2
PSI = 0.077
Binding site
SNORD22
PRPF8
EFTUD2
NMD-targeted transcript
PTC
UG
A
Protein-coding transcript
100 vertebrates conservation by PhastCons
B
PSI (%)
NC5
ASO1
ASO2
SNORD22 KD
C
Splice site score
8.95
3.88
9.06
11.18
Strong
Moderate
Weak
SRRT 5'
3'
D
Proximal to downstream 5'SS
Distal to downstream 5'SS
SNORD22
SRRT 5'
3'
MTM-2

**Figure S20. The SNORD22–PRPF8–EFTUD2 complex promotes inclusion of an NMD-associated cassette exon in *SRRT*, related to Figure 5A**. **(A)** Sashimi plots of *SRRT* showing the cassette exon event identified by rMATS upon SNORD22 KD. The negative control (NC5, two replicates) is shown in black and the SNORD22 KD condition (two ASOs, two replicates each) is shown in blue. The cassette exon is highlighted in gray and PSI values for this exon in each sample are indicated above its position. Binding sites of SNORD22, PRPF8 and EFTUD2 are shown below the sashimi plots along with two *SRRT* transcript isoforms: the upper transcript contains the cassette exon, introducing a premature stop codon (NMD-targeted isoform) whereas the lower transcript lacks the cassette exon and corresponds to the canonical SRRT isoform. The PhastCons 100-vertebrate conservation track is shown in green with scores ranging from 0 to 1 (higher values indicate higher conservation). **(B)** ddPCR validation of exon exclusion upon SNORD22 depletion. Bars represent mean ± SE and points represent individual biological replicates ($n$ = 3). Statistical significance was assessed using two-sided $t$-test comparing NC5 to each SNORD22 ASO (*$p < 0.05$). **(C)** Bar plot of splice site strength scores for the upstream and downstream introns flanking the cassette exon. Splice site strength was computed using MaxEntScan and categorized as weak (score < 8), moderate (8 ≤ score ≤ 9) or strong (score > 9). **(D)** Base-pairing interactions between SNORD22 and two binding sites in *SRRT* corresponding to the regions shown in (A). The MTM-2 motif identified in Figure 3E is highlighted within the SRRT sequence.

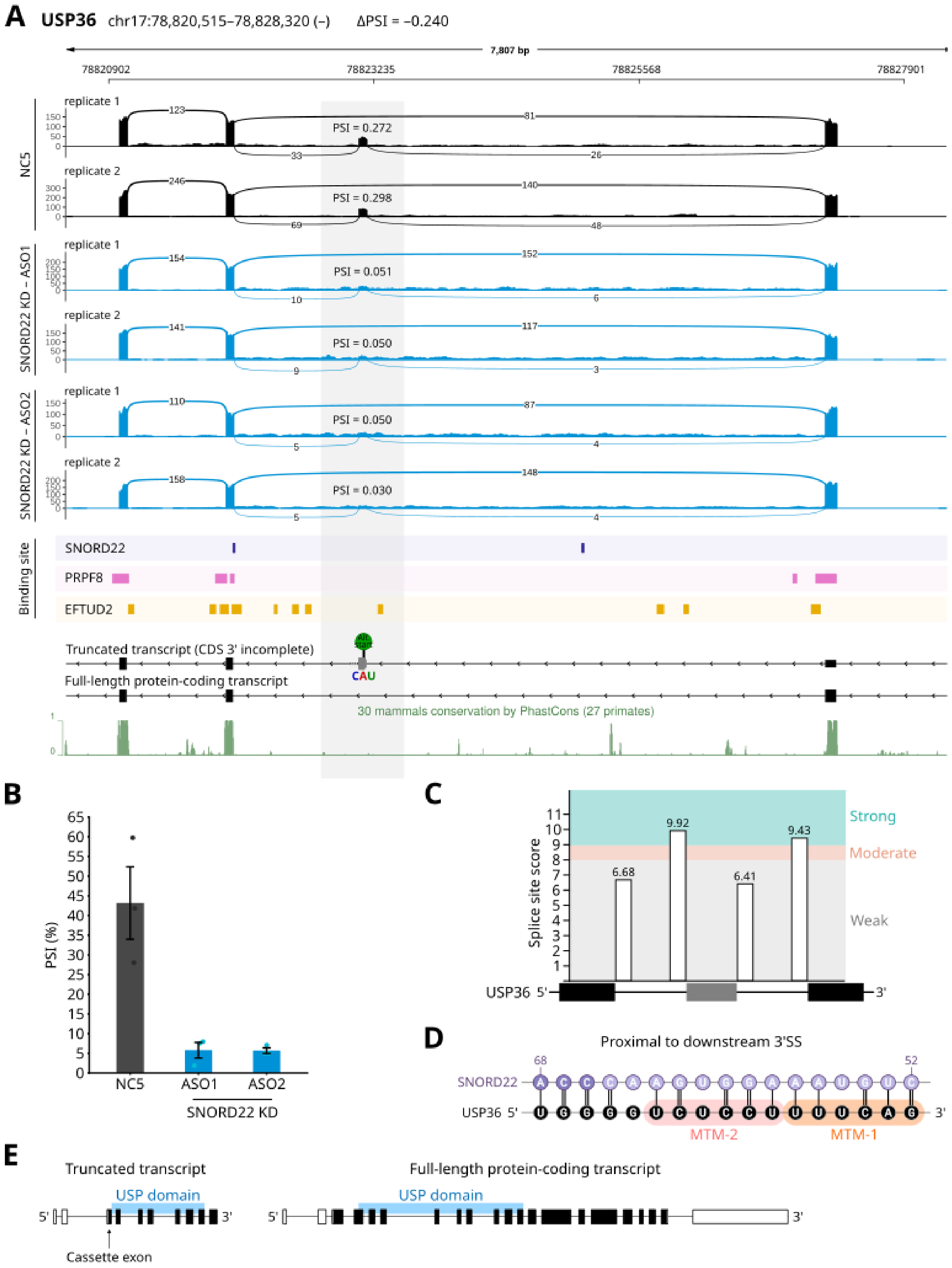
A
USP36 chr17:78,820,515–78,828,320 (–) ΔPSI = –0.240
7,807 bp
78820902
78823235
78825568
78827901
NC5
replicate 1
PSI = 0.272
replicate 2
PSI = 0.298
SNORD22 KD – ASO1
replicate 1
PSI = 0.051
replicate 2
PSI = 0.050
SNORD22 KD – ASO2
replicate 1
PSI = 0.050
replicate 2
PSI = 0.030
Binding site
SNORD22
PRPF8
EFTUD2
Truncated transcript (CDS 3' incomplete)
CAU
Full-length protein-coding transcript
30 mammals conservation by PhastCons (27 primates)
B
PSI (%)
NC5
ASO1
ASO2
SNORD22 KD
C
Splice site score
6.68
9.92
6.41
9.43
Strong
Moderate
Weak
USP36 5'
3'
D
Proximal to downstream 3'SS
SNORD22
USP36 5'
3'
MTM-2
MTM-1
E
Truncated transcript
USP domain
Cassette exon
Full-length protein-coding transcript
USP domain

**Figure S21. The SNORD22–PRPF8–EFTUD2 complex is associated with a cassette exon in *USP36* that defines a truncated isoform, related to Figure 5B. (A)** Sashimi plots of *USP36*, shown in the same format and color scheme as in Figure S20A. The cassette exon-containing transcript uses an alternative start codon within the cassette exon and is annotated in Ensembl as a CDS 3′ incomplete (truncated) isoform. The cassette-exon skipping transcript corresponds to the full-length protein-coding USP36 isoform. The PhastCons 30-mammal conservation track is shown below. **(B)** ddPCR validation of exon exclusion upon SNORD22 depletion. Bars represent mean ± SE and points represent individual biological replicates ($n$ = 3). Statistical significance was assessed using two-sided $t$-test comparing NC5 to each SNORD22 ASO ($p = 0.052$, $p = 0.054$). **(C)** Bar plot of splice site strength scores for the upstream and downstream introns flanking the cassette exon, shown in the same format and color scheme as in Figure S20B but for USP36. **(D)** Base-pairing interactions between SNORD22 and the binding site on USP36 proximal to the downstream 3' splice site. MTM-1 and MTM-2 motifs introduced in Figure 3E are highlighted within the USP36 sequence. **(E)** Schematic comparing the full-length USP36 transcript (cassette exon absent) and the truncated transcript (cassette exon included). The USP catalytic domain is annotated for both transcripts.
